# The *de novo* molecular evolution of a viper venom metalloproteinase inhibitor and its target network

**DOI:** 10.64898/2026.09.24.752146

**Authors:** Noah L. Dowell, Susana Vázquez Torres, Fiona P. Ukken, Milorad Andjelkovic, Mamta Hajra, Elizabeth Cahill, Birte Höcker, Rafael Fernández-Leiro, Sean B. Carroll

**Author notes:** Co-first authors (Noah L. Dowell, Susana Vázquez Torres). Corresponding author: Sean B Carroll.

## Abstract

Molecular innovation and coevolution play major roles in the evolution of biological complexity and diversity, yet we have few instances in which we understand how proteins acquired novel biological and biochemical functions. Rattlesnakes and other vipers have evolved protection against their own dangerous venom metalloproteinase (MP) toxins via the invention and coevolution of a novel class of serum-borne MP inhibitors (FETUAs) derived from an ancestral, non-inhibitory Fetuin-A glycoprotein. However, how FETUAs acquired their function as well as their mechanism of action are unknown. Here, we use a combination of structure prediction, rational mutagenesis, functional analyses, and molecular dynamics to elucidate how the rattlesnake FETUA-3 protein evolved to inhibit three classes of venom MP toxins. We show that numerous changes across three structural elements were required for FETUA-3 to bind to multiple MP targets with high-affinity (∼nM), including substitutions in a short N-terminal region that enabled it to insert into MP active sites and directly disrupt substrate cleavage. We also find strong functional and evolutionary constraints on key N-terminal residues that are required to maintain FETUA-3’s multi-target activity. Our results reveal that FETUA-3 convergently evolved a unique mechanism of action distinct from other vertebrate metalloproteinase inhibitors. We suggest that the scope of molecular evolution of the FETUA-3 protein reflects that in general, as compared to shifts in substrate or ligand specificity, a longer evolutionary path is necessary for the de novo evolution of high affinity protein-protein interactions and networks.

## Introduction

Key innovations are a distinct category of novel traits that enable organisms to adapt to new environments and lifestyles and can catalyze adaptive radiations (1). The advent of such traits can in turn spur coevolutionary adaptations both within lineages that bear them and in species with which they interact (2–4). Understanding the mechanistic basis of innovation and coevolution, then, are major aims of evolutionary biology (5,6) which requires the tracing of the molecular origins of key innovations and their coevolutionary adaptations.

One central question in the evolution of biochemical innovations concerns how proteins acquire new specific interactions and functions. A growing body of work has suggested that functional shifts in ligand-binding or substrate specificity may require only a small number of large-effect mutations (7) and thus may be relatively simple to evolve. However, most of these examples are not linked to key innovations, and they generally entail a small number of protein-contacts and not the much larger interfaces characteristic of protein-protein complexes (8).

A second general question regarding the evolution of protein innovation concerns how many molecular “solutions” may exist for a new biological function, and to what degree a novel protein function comprises an entirely new solution or the exploitation of a pre-existing activity. The frequent occurrence of convergent evolution in which different organisms acquire similar traits independently suggests that at the phenotypic level, a limited set of solutions is available or favored. At the molecular level, functional convergence spans a spectrum from identical substitutions in the same protein molecules in multiple disparate species (9) to the origin of the same biochemical function or catalytic activity by entirely unrelated proteins in different kingdoms (see refs.10,11). Because the origins of few key biochemical innovations have been elucidated, it is not clear in general to what degree their functions have emerged *de novo* or from pre-existing activities.

Venomous animals are rich models for exploring these questions because their venoms are key protein innovations that provided new means of capturing prey, arose independently in multiple lineages, and spur coevolutionary adaptations(12). Snake venoms, for example, typically contain an arsenal of toxins belonging to 10-15 protein families(13,14) which can drive the evolution of venom resistance in prey(15). Prey resistance can, in turn, select for changes in snake venom composition(16). The snake-prey interspecific arms race is also accompanied by key intraspecific coevolutionary adaptations within the venomous snakes themselves. Most snake venoms disrupt hemostasis or neural transmission by acting on molecular targets that are deeply conserved among vertebrates, including snakes. Therefore, many groups of snakes have coevolved auto-resistance to their own venoms either through the coevolution of changes in endogenous toxin receptors in neurotoxic snakes (17,18) or through the coevolution of circulating venom toxin inhibitors in hemotoxic snakes (19).

For instance, in vipers (Viperidae), one of the two largest families of venomous snakes that originated approximately 50 million years ago(20), the metalloproteinase (MP) family is often the most abundant and structurally diverse family of toxins (21,22). The MP toxin family arose in snakes from the vertebrate *ADAM28* metalloproteinase gene(22,23), a member of the large ADAM (<u>a</u> <u>d</u>isintegrin <u>a</u>nd <u>m</u>etalloproteinase) family within the metzincin superfamily of zinc-dependent proteases. During the evolution of vipers, the MP toxin family has expanded and diversified into three structural classes (24) that attack broadly conserved extracellular matrix, basement membrane, and blood clotting proteins causing hemorrhage, organ damage, and coagulopathy in their prey or accidental human victims(25). To protect themselves against these dangerous toxins, vipers did not exploit existing MP/ADAM inhibitors such as the deeply conserved and broad-acting tissue inhibitor of metalloproteinase (TIMP) family(26,27) or the astacin-type MP inhibitor Fetuin-B(28). Rather, the snakes coevolved a novel class of serum-borne MP auto-inhibitors from the non-inhibitory, ancestral vertebrate serum glycoprotein Fetuin-A, we refer to these as FETUA proteins(29,30). There are four FETUA proteins in *Crotalus atrox* that collectively inhibit the enzymatic, hemorrhagic, and lethal activities of its ∼30 venom MPs(29,30), and together constitute a key innovation that enables the venomous lifestyle.

Understanding the mechanism of FETUA action is of dual interest. First, because these novel inhibitors evolved from a non-inhibitory ancestral protein, it is of fundamental interest to elucidate how FETUAs acquired their function and to compare their mode of action with other vertebrate MP inhibitors. And second, because these proteins have shown notable promise as antivenom agents *in vivo*(30), it is important to understand how a small set of FETUA proteins controls the activity of a large, diverse set of MPs. Here, we use a combination of structure prediction, rational mutagenesis, functional assays, and molecular dynamics to identify multiple molecular determinants that enable the FETUA-3 protein to bind to a diverse subset of venom MPs with high affinity, and to insert its unique N-terminal domain into MP enzyme active sites and inhibit substrate cleavage. FETUA-3’s mechanism of action is distinct from other classes of MP inhibitors and thus constitutes a case of functional convergence. We propose that the extent of the molecular evolution of the FETUA-3 protein reflects that, in general, more numerous intermolecular contacts must evolve to acquire new high affinity protein-protein interactions than to shift protein specificity for small molecule ligands or substrates.

## Results

### FETUA-3 is a high-affinity, multi-target, non-competitive metalloproteinase inhibitor

We have previously shown that the *C. atrox* FETUA-3 protein inhibits the activity of three of the most abundant *C. atrox* venom metalloproteinases belonging to three different MP structural classes(29) that differ by the presence/absence of three distinct domains: MDC-4 (48 kDa), a class III MDC toxin possessing a metalloproteinase, disintegrin-like and cysteine-rich domain; MPO-1 (∼23 kDa), a class I MP class possessing only the metalloproteinase domain; and MAD-3 (∼23 kDa), a class II MP also consisting of only a metalloproteinase domain due to post-translational cleavage of the disintegrin domain in the venom gland (Figure 1A). Our MAD-3 preparation contains two closely related paralogs, MAD-3a and MAD-3b, that are 90% identical across the metalloproteinase domain and enzymatically equivalent for the purposes of this study (see Material and Methods).

**Figure 1.**
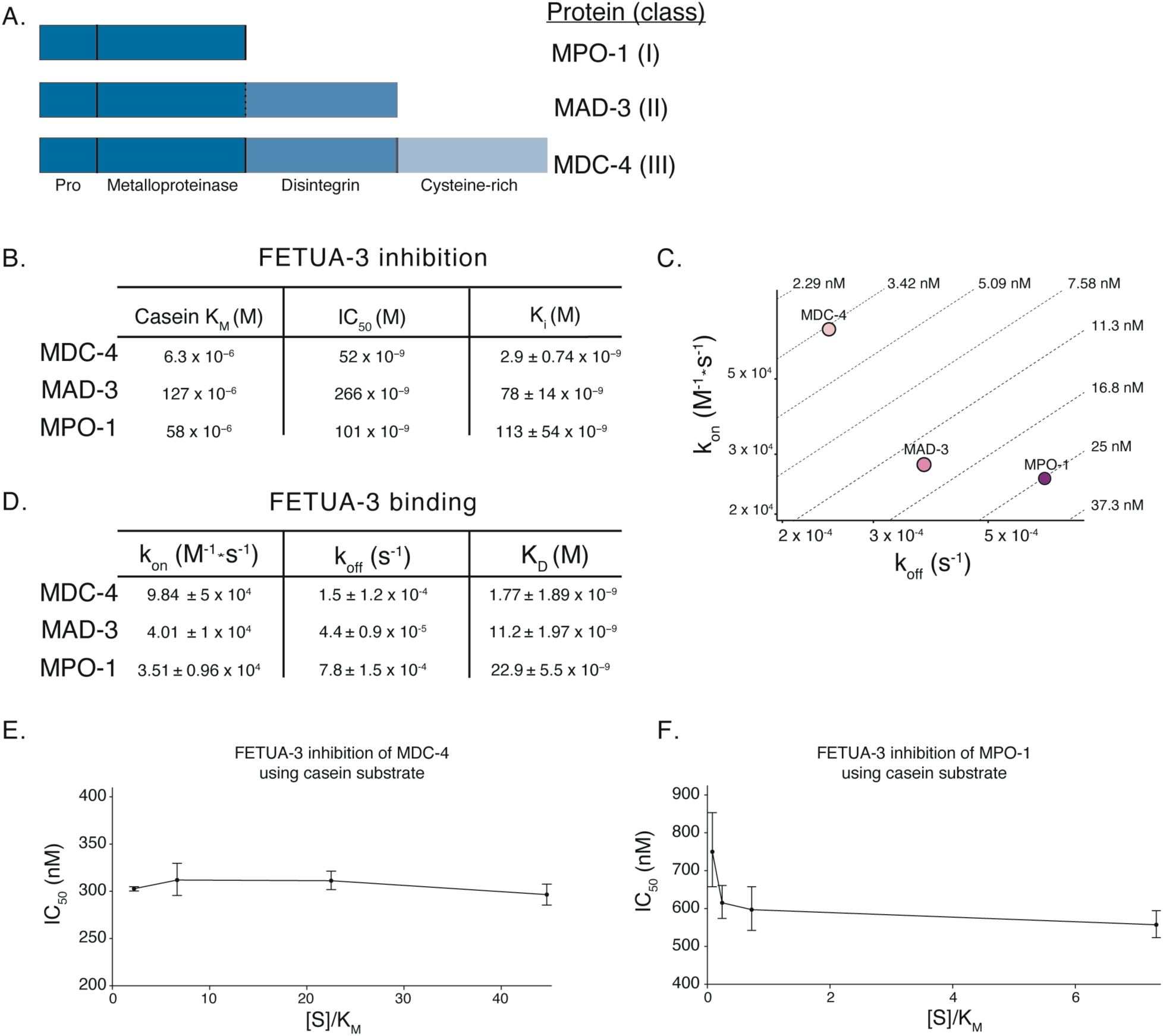
FETUA-3 is a high affinity, multi-target, non-competitive venom metalloproteinase inhibitor. (A) Schematic diagram of the protein domain organization for MPO-1, MAD-3 and MDC-4. The abbreviation “Pro” represents the pro-domain which is absent from the venom secreted metalloproteinases. The dashed line between the metalloproteinase and disintegrin domains of MAD-3 represents the proteolytic cleavage site. The diagram is not drawn to scale. (B) FETUA-3 strongly inhibits MDC-4, MAD-3, and MPO-1. The Michaelis-Menten constant (K_M_) for MDC-4, MAD-3 and MPO-1 cleavage of casein shows the variation in catalytic activity on a common substrate. Titration of FETUA-3 inhibition of casein cleavage by metalloproteinases was used to establish the inhibitor concentration at which half of the enzyme is inhibited (IC_50_). This measure of inhibition is specific to the experimental conditions for each enzyme and not comparable across enzymes. FETUA-3 inhibition constants (K_i_ ± standard deviation) for all three enzymes account for differences in cleavage efficiency of casein and allow for the comparison of inhibitor potency across enzymes. (C) FETUA-3 binds to MDC-4 more strongly than to MAD-3 or MPO-1. The iso-affinity plot of a representative experiment shows the association rate constant (k_on_ (M^-1^*s^-1^); y-axis) versus the dissociation rate constant (k_off_ (s^-1^); x-axis) with the diagonal representing the equilibrium dissociation constant (K_D_ (M); dashed diagonal lines). The lower binding affinities of FETUA-3 with MAD-3 and MPO-1 relative to MDC-4 are due their slower association and faster dissociation rates constants. (D) The association (k_on_) and dissociation rate constants (k_off_) and equilibrium dissociation constants (K_D_ ± standard deviation, n = 3) for interactions between FETUA-3 and MDC-4, MAD-3 and MPO-1. (E) FETUA-3 is a non-competitive inhibitor of MDC-4. Replot (IC_50_ vs [casein]/K_M_) of FETUA-3 inhibition of MDC-4 across a range of casein (2.2 – 44.7 x K_M_) concentrations shows that greater substrate concentrations do not affect the IC_50_, indicating non-competitive inhibition. (F) FETUA-3 is a non-competitive inhibitor of MPO-1. Replot (IC_50_ vs [casein]/K_M_) of FETUA-3 inhibition of MPO-1 across a range of casein (0.08 – 7.3 x K_M_) concentrations shows that greater substrate concentrations do not affect the IC_50_, indicating non-competitive inhibition.

In order to quantify and compare the strength of FETUA-3 inhibition of enzymatic activity *in vitro* for all three MPs, we selected succinylated casein as a substrate because MPs exhibit different substrate preferences (e.g., MAD-3 does not cleave collagen I), and succinyl casein is a common substrate for all three enzymes. To determine the enzyme inhibition constants of FETUA-3, we first determined the Michaelis-Menten constants (K_M;_ 1/2V_max_) for each enzyme on succinyl casein (“casein” hereafter) and found that MDC-4 had the highest affinity (6.3 x 10^-6^ M), MPO-1 had moderate affinity (58 x 10^-6^ M) and MAD-3 had the lowest affinity (127 x 10^-6^ M) (Figures 1A and S1). We then titrated FETUA-3 inhibition against all three MPs and obtained inhibition constants (K_i_) for MDC-4 (K_i_ = 2.9 ± 0.74 x 10^-9^ M), MAD-3 (K_i_ = 78 ± 14 x 10^-9^ M) and MPO-1 (K_i_ = 113 ± 54 x 10^-9^ M), (Figure 1A and see Materials and Methods).

In order to determine how enzyme inhibition may be related to FETUA-3 binding to these MPs, we used surface plasma resonance (SPR) to measure the kinetics of interactions between FETUA-3 and each of the three venom MPs. FETUA-3 exhibited the highest affinity for MDC-4 (K_D_ = 1.8 ± 1.89 x 10^-9^ M) and strong affinity for MAD-3 (K_D_ = 11.2 ± 1.97 x 10^-9^ M) and MPO-1 (K_D_ = 22.9 ± 5.5 x 10^-9^ M) (Figures 1B and 1C). Thus, FETUA-3 is a high affinity and potent inhibitor of the three target toxins.

Since FETUA-3 is a novel MP inhibitor, there is no basis for predicting the mode of inhibition of MP activity (i.e. competitive or non-competitive). To investigate how FETUA-3 behaves, we measured the strength of inhibition (IC_50_) while varying the substrate concentration relative to the enzyme-specific K_M_ and plotted the relationship between IC_50_ and substrate concentration(31). We found that FETUA-3 inhibited MDC-4 and MPO-1 across a wide concentration range of casein (MDC-4: 2.2 – 44.7 x K_M_ and MPO-1: 0.08 – 7.3 x K_M_) and observed minimal change in the strength of inhibition (Figures 1G, H). These data indicate that FETUA-3 is a noncompetitive inhibitor of MDC-4 and MPO-1. This mode of inhibition appears to make biological sense in that FETUA-3 functions in protein-rich environments (blood, extracellular space) where metalloproteinase substrates such as clotting components and extracellular matrix proteins are present in excess relative to the inhibitor concentration.

To elucidate how FETUA-3 evolved to act as a high-affinity, multi-target, non-competitive inhibitor, we sought to identify FETUA-3 protein regions that may be involved in MP binding and inhibition.

### Structure prediction reveals several candidate FETUA-3 regions that engage venom metalloproteinases

To explore how FETUA-3 may interact with multiple venom MPs, we used a variety of structure prediction methods. Deep learning-based structure-prediction methods have recently advanced, particularly with respect to predicting biomolecular interactions(32). Moreover, high resolution structures of a large number of snake venom metalloproteinases have been determined(33) including MDC-4(34) (Vap2b) and Atrolysin-C(35) (a very close homolog of MPO-1), and Fetuin-A (the ancestor of FETUA proteins) is a member of the well-studied cystatin protein superfamily(36).

We first used AlphaFold3(AF3)(32) to predict structures of FETUA-3 in complex with the metalloproteinase domains of MDC-4, MAD-3b and the MPO-1 enzyme because the metalloproteinase domain is the only domain shared by these enzymes (the pairwise sequence identity of these three ∼ 208 amino acid long metalloproteinase domains ranges from 51-63%) . This approach allows direct comparison across MP classes, although it does not capture contributions from the disintegrin-like and cysteine-rich domains of class II and III enzymes, which in intact proteins could influence interface accessibility or provide additional contacts.

We generated 125 models per complex from 25 independent seeds, each producing five diffusion samples, and selected a highly ranked model for analysis (Figure 2A, selected models corresponding ipTM and pTM values in top panel; ranks of selected models identified with black triangles in lower panel). The range of Interface Predicted Template Modeling (ipTM) and Predicted Template Modeling (pTM) confidence scores differed sharply between complexes (Figure 2A, lower panel and Figure S2A). In aggregate, FETUA-3 complexed with MAD-3b was predicted confidently (ipTM 0.86 ± 0.06, pTM 0.82 ± 0.04), whereas complexes with MDC-4 (ipTM 0.39 ± 0.18, pTM 0.61 ± 0.08) and MPO-1 (ipTM 0.42 ± 0.16, pTM 0.59 ± 0.06) were not (Figure 2A, average ipTM and pTM values in box below x-axis in lower panel).

**Figure 2.**
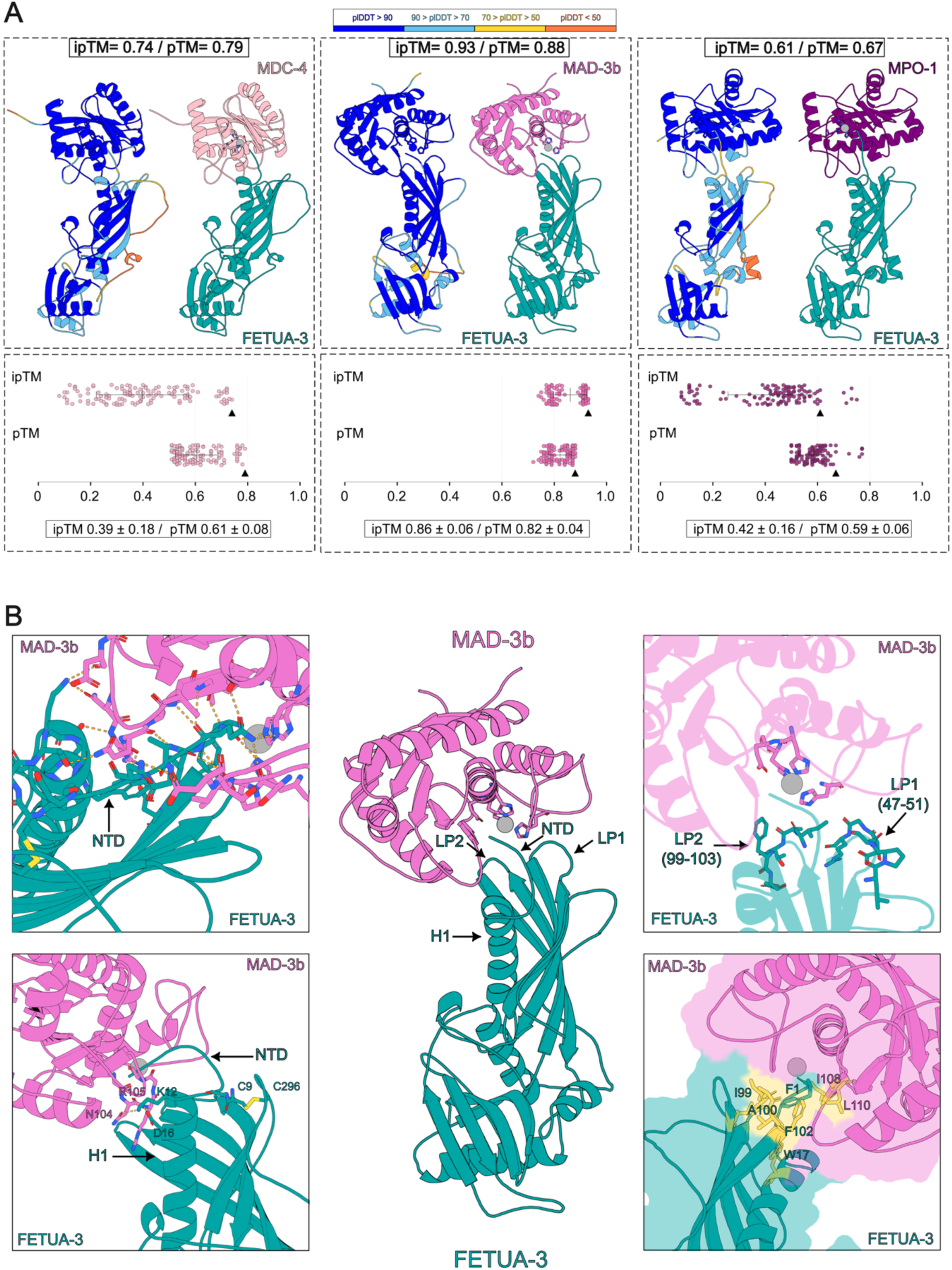
AlphaFold3 modeling of FETUA-3-MP complexes predicts four interaction interfaces. (A) Top: AlphaFold3 models of FETUA-3 bound to the metalloproteinase domains of MDC-4 (class III, left), MAD-3b (class II, center) and MPO-1 (class I, right). Within each panel, on the left, each complex is colored according to predicted local distance difference test (pLDDT) confidence scores and by chain while on the right FETUA-3 is in teal and each metalloproteinase in the indicated color; the catalytic zinc is a gray sphere. Predicted interface TM-score (ipTM) and predicted TM-score (pTM) values for the displayed model are given above each panel. The displayed models are the highest-ranked models for each complex, except for MPO1 which is the fourth-ranked model (the highest ranked model placed FETUA-3 in an orientation inconsistent with the FETUA-3-MAD-3b complex) Bottom: ipTM (upper row) and pTM (lower row) across all 125 models per complex (25 seeds × 5 samples). Each point is one model, black bars are mean ± SD (values below), black triangles mark the displayed model, and dashed lines indicate the 0.6 and 0.8 confidence thresholds. (B) Predicted structure of the FETUA-3–MAD-3b complex. Top left, the N-terminal domain of FETUA-3 forms extensive electrostatic interactions with regions of MAD-3b both proximal and distal to the active site. Top right, loop 1 (LP1) and loop 2 (LP2) are positioned adjacent to the enzyme active site. Bottom left, a disulfide bond between Cys9 and Cys296 of FETUA-3 is predicted to stabilize the N-terminal domain in a conformation compatible with inhibition. Bottom right, the N-terminal residue Phe1 (F1) of FETUA-3 inserts into a hydrophobic pocket formed by LP2 residues Ile99, Ala100 and Phe102 of the inhibitor together with enzyme residues Ile108 and Leu110.

To test whether these differences reflect a limitation of one method, we analyzed the same complexes with ESMFold2(37) and OpenDDE(38). For FETUA-3 complexed with MAD-3b, all three methods were consistent in returning prediction scores with high confidence (range of ipTM and pTM 0.8 to 1.0; average scores of 125 models for three methods: ESMFold2 ipTM 0.92 ± 0.09, pTM 0.88 ± 0.05; OpenDDE ipTM 0.90 ± 0.03, pTM 0.87 ± 0.02) (Figure S2B – C and Figure S3A, B). For the other two predicted complexes, the average scores were below the low confidence threshold (ipTM or pTM < 0.6; see Figure S2 for average scores), while the highest-scoring AF3 and ESMFold-2 predictions for FETUA-3 and MDC-4 and MPO1 fell between low (0.6) and high (0.8) confidence thresholds (zone of uncertainty).

Low interface confidence could in principle reflect genuinely weaker FETUA3-MP binding rather than prediction failure. However, our experimental data exclude this possibility, since FETUA-3 binds and inhibits MDC-4 and MPO-1 with higher affinity and potency than those measured for MAD-3b (Figure 1). Thus, the lower ipTM values for MDC-4 and MPO-1 reflect a limitation of the prediction.

Since all three MPs are high affinity targets, our initial strategy was to set aside the lower confidence models for the moment and to utilize the highest confidence FETUA-3-MAD-3b model for inferences of potential interacting regions (Figure 2B), then to conduct experiments to assess the functional contributions of these features to the binding and inhibition of all three targets. We will later revisit the lower confidence models in light of additional functional data.

The majority of the FETUA-3 protein (which includes two cystatin domains) is confidently predicted (Predicted Local Distance Difference Test (pLDDT) (Figure 2A, ribbon representations on left side of panels with pLDDT > 90 shown in dark blue). Lower confidence is confined to a discrete disordered region distant from the target enzyme (Figure 2A, pLDDT < 50 to 70 shown in orange and yellow) whereas the elements at the FETUA-3 and metalloproteinase interface are confidently predicted. Four FETUA-3 structural domains point towards the interface with MAD-3b (Figure 2B, black arrows in central panel). Starting from the N-terminus these four domains are: i) the N-terminal domain (“NTD”) (Figure 2B, top left); ii) the first alpha helix (“helix 1 or H1”) (Figure 2B, bottom left); iii) the first hairpin loop (“loop 1 or LP1”) (Figure 2B, top right); and iv) the second hairpin loop (“loop 2 or LP2”) (Figure 2B, top right)). In addition, the model places cysteine-9 (C9) and cysteine-296 (C296) in positions compatible with disulfide bond formation that may orient the NTD toward the MP active site (Figure 2B, lower left). Prior studies of the ancestral protein FETUIN-A have identified a disulfide bridge between the highly conserved first (C9) and twelfth (C296) cysteines(39).

Each element contributes distinct predicted contacts. The NTD forms extensive electrostatic interactions with the enzyme that could stabilize the complex (Figure 2B, top left, orange dashed lines linking oxygen and nitrogen atoms). Phe1 of FETUA-3 inserts into a hydrophobic pocket formed at the interface by residues from both chains, comprising I99, A100 and F102 of loop 2 together with A108 and L110 of MAD-3b (Figure 2B, lower right, hydrophobic surface in yellow). Helix 1 (Figure 2B, lower left) and loop 1 (Figure 2B, upper right) contribute further electrostatic contacts, suggesting that these regions may also participate in target recognition. Overall, 24 FETUA-3 interface residues are predicted to interact with 23 MAD-3b residues.

To test these predictions that five structural elements of FETUA-3, including the NTD, a disulfide bond, part of an alpha helix and two hairpin loops may be involved in MP inhibition and binding, we produced and analyzed the activities of recombinant FETUA-3 proteins in which each of these elements is altered.

### The FETUA-3 N-terminal domain sequence and spatial position are necessary for MP inhibition and binding

To test the prediction that NTD interacts with MPs near the active site, we made a recombinant FETUA-3 protein that lacks the first seven amino acids of the protein (FETUA-3-delNTD, Figure 3A); we did not remove the eighth amino acid (D8) because that would generate a protein with an amino-terminal cysteine residue (C9) that might not fold properly or could be less stable. We found that the inhibition of MAD-3 activity was abolished in the FETUA-3-delNTD protein (Figure 3B) as well as the inhibition of MDC-4 and MPO-1 activity (Figures 3C, D), revealing that the first seven amino acids are necessary for FETUA-3 function on all three MPs.

**Figure 3.**
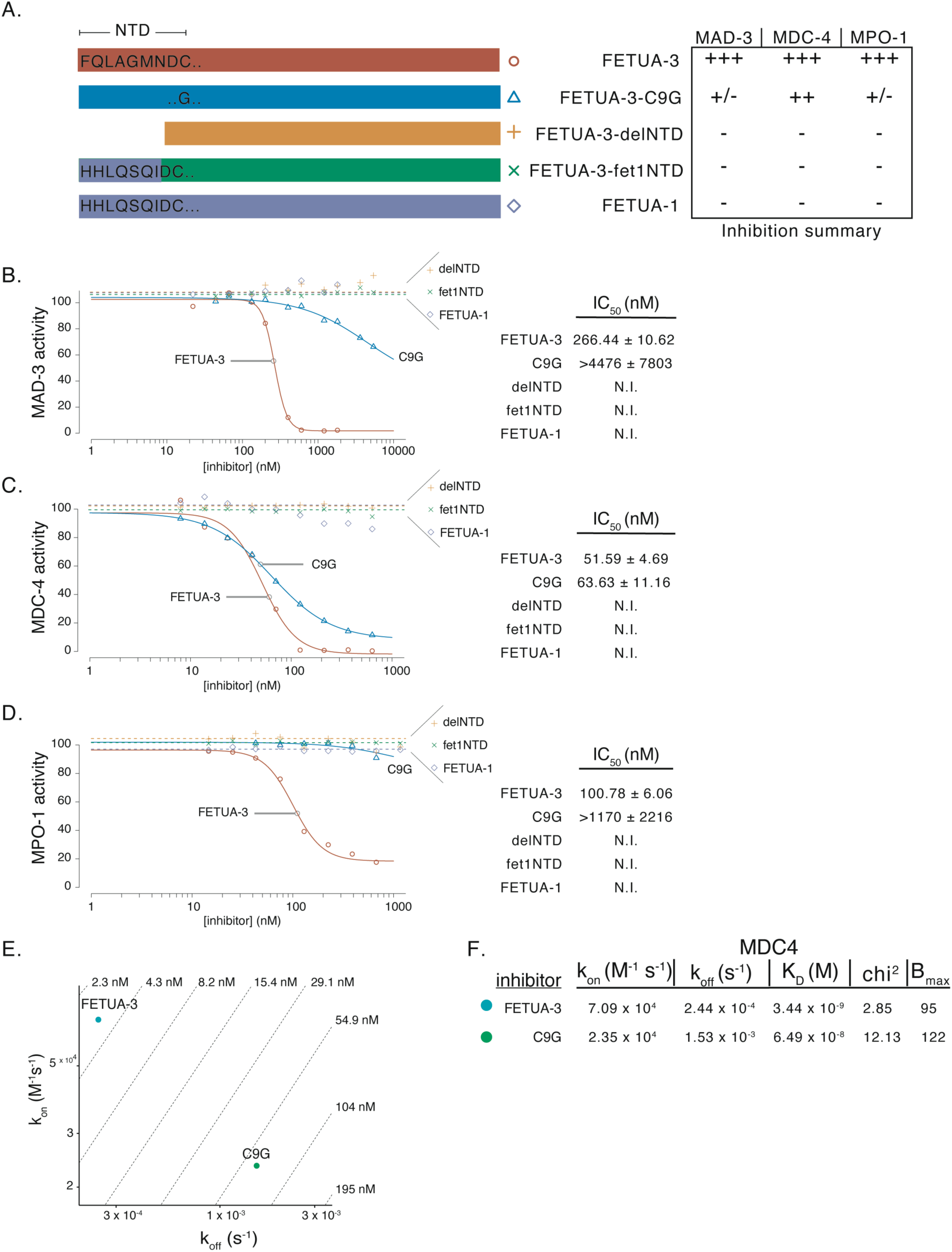
The FETUA-3 N-terminal domain sequence and spatial position are necessary for MP inhibition and binding. (A) A schematic representation of the mutant or chimeric FETUA-3 proteins tested for inhibition of and binding to MDC-4, MAD-3 and MPO-1. Top, the N-terminal domain amino acid sequence (1-FQLAGMNDC-9) of FETUA-3 after removal of the signal peptide (red rectangle). Second row, a single substitution at position nine of a cysteine for a glycine (FETUA-3-C9G) (blue rectangle). Third row, the FETUA-3-delNTD protein (orange rectangle) lacks the first seven amino acids and begins with an N-terminal aspartic acid (D8). Fourth row, the chimeric FETUA-3-fet1NTD (green rectangle) consists of a swap of the first seven amino acids of FETUA-3 to the FETUA-1 sequence (1-HHLQSQI-7) while the rest of the protein is FETUA-3 sequence. Fifth row, FETUA-1 (purple rectangle), a non-inhibitor of venom metalloproteinases, provides donor sequences that are likely functional as a FETUA sequence but lacks binding or inhibitory activity with respect to MDC-4, MAD-3 and MPO-1. A summary of the inhibition data presented in B – D is shown to the left of the protein schematics and uses the protein names to label the rows of the table. (B-D) Inhibition of MAD-3 (B), MDC-4 (C) and MPO-1 (D) by FETUA-3, FETUA-3-C9G (C9G), FETUA-3-delNTD (delNTD), FETUA-3-fet1NTD (fet1NTD) and FETUA-1. The colors of the lines are matched to the rectangles in (A) and the midpoints (IC_50_ ± standard error, nM) of the curves are reported in the tables on the right. N.I.= no inhibition is detected. (B) Deletion or replacement of the N-terminal domain abolishes, and the C9G substitution very strongly reduces FETUA-3 inhibition of MAD-3. (C) Deletion or replacement of the N-terminal domain abolishes FETUA-3 inhibition of MDC-4, but the C9G substitution has only a slight effect. (D) Deletion or replacement of the N-terminal domain, and the C9G substitution abolish FETUA-3 inhibition of MPO-1. (E) An iso-affinity plot of a representative experiment shows that the ∼19-fold reduction of the FETUA-3-C9G equilibrium dissociation constant from MDC-4 is due to a slower on-rate (k_on_) and a faster off-rate (k_off_). (F) A table showing association and dissociation rate constants and equilibrium dissociation constant for MDC-4 interactions with FETUA-3 and FETUA-3-C9G.

The removal of the seven N-terminal amino acids of the FETUA-3 protein could disrupt the protein’s structure in some way. To distinguish whether it is the sequence of these seven amino acids that is important for FETUA-3 function, or merely the presence of seven amino acids, we also made a construct in which the first seven amino acids were replaced with the first seven amino acids of the native FETUA-1 protein, the liver expressed, non-inhibitory FETUA protein orthologous to vertebrate FETUIN-A from which inhibitory FETUAs evolved (FETUA-3-fet1NTD, Figure 3A). We also did not detect any inhibition of MAD-3, MDC-4, or MPO-1 activity by the FETUA-3-fet1NTD protein (Figures 3B-D).

The complete loss of MP inhibition by deletion or replacement of the first seven amino acids of the N-terminal region/domain of FETUA-3 is consistent with the FETUA-3-MAD-3b model that suggests that the N-terminal domain inserts into the MP active site. These data lead us to consider two scenarios for how the NTD could inhibit MP activity. One possibility is that the NTD mediates target recognition and thus is key for binding to MPs. Alternatively, the NTD may inhibit substrate cleavage by disruption of the active site after binding. If the latter is the case, we might expect to observe MP binding by the FETUA-3-delNTD and FETUA-3-fet1NTD proteins, while in the former scenario we would not expect to observe MP binding. We measured binding interactions between the three venom MPs and FETUA-3-delNTD, FETUA-3-fet1NTD, FETUA-1 and wild-type FETUA-3 in single cycle kinetic experiments. We did not detect any binding to MPs by FETUA-3 proteins in which the NTD is deleted (Figures S4A - F) or replaced by the FETUA-1 NTD (FETUA-1 does not bind the three venom MPs we tested; Figure S4A - F). These results indicate that the NTD of FETUA-3 is necessary for both binding and inhibition of the three MP enzymes, and that the NTD is one interface that evolved in the origin of the FETUA-3 inhibitor from the ancestral FETUA-1 protein.

Prior studies of FETUIN-A proteins have shown that C9 forms a disulfide bond with C296(39). In the AlphaFold model of FETUA-3-MAD-3b, C9 and C296 are within 3 angstroms (Å), which supports the inference of a disulfide bond at this position that would bring the C-terminal region of FETUA-3 very close to the metalloproteinase and may contribute to the positioning of the NTD (aa 1-7) in an inhibitory pose within the MP active site. To test this possibility, we made recombinant FETUA-3 with a cysteine to glycine substitution at residue nine (FETUA-3-C9G, Figure 3A) to abolish the disulfide bond and measured the ability of this mutant protein to inhibit and bind to the three MPs.

We observed that the C9G mutation strongly reduced inhibition of MAD-3 (Figure 3B, (IC_50_ > 4476 nM)) and of MPO-1 (Figure 3D, IC_50_ > 1170 nM), whereas we observed a modest, 1.2-fold increase in the IC_50_ of FETUA3-C9G (64 nM) versus wild-type FETUA-3 (52 nM) against the MDC-4 enzyme (Figure 3C). The C9G mutation’s effects on FETUA-3 binding affinities for the respective MPs are similar to its effect on enzyme inhibition: binding to MDC-4 is reduced about 19-fold (Figure 3E, F and Figure S5A; C9G binding to MDC-4 suggests a properly folded mutant protein) and binding to MAD-3 (Figure S5B) and MPO-1 (Figure S5C) is not detected.

Together, these results suggest that the FETUA-3 N-terminal domain is necessary for MP inhibition and binding and it is positioned in the MP active site in an inhibitory pose by a disulfide bond between C9 and C296. We were surprised that a single amino acid substitution disrupting a disulfide bond resulted in the loss of binding and inhibition for two enzymes (MAD-3 and MPO-1) but only a slight decrease of binding and inhibition for the third enzyme (MDC-4). This raises the possibility that FETUA-3 functions as a multi-target inhibitor through additional regions of the protein which may vary in their contribution to the binding and inhibition of different MPs. Therefore, we next tested whether the other identified FETUA-3 domains are also important for target binding and inhibition.

### Two additional FETUA domains are required for multi-target inhibition

Three candidate regions outside of the NTD may interact with and contribute to the inhibition of venom MPs by FETUA-3 including: i) a portion of the first alpha helix (“helix 1, H1”); ii) a hairpin loop between beta-strands, corresponding to amino acids 47-PSDGR-51 (“loop 1, LP1”); and iii) a second hairpin loop 99-IATFE-103 (“loop 2, LP2”) (Figure 4A). These structural elements are also present in the model of the non-inhibitor FETUA-1. Therefore, we tested whether specific FETUA-3 regions are necessary for MP inhibition or binding by replacing the FETUA-3 sequences with the homologous (and well-conserved) FETUA-1 sequences to make FETUA-3-fet1 chimeric proteins (Figure 4A).

**Figure 4.**
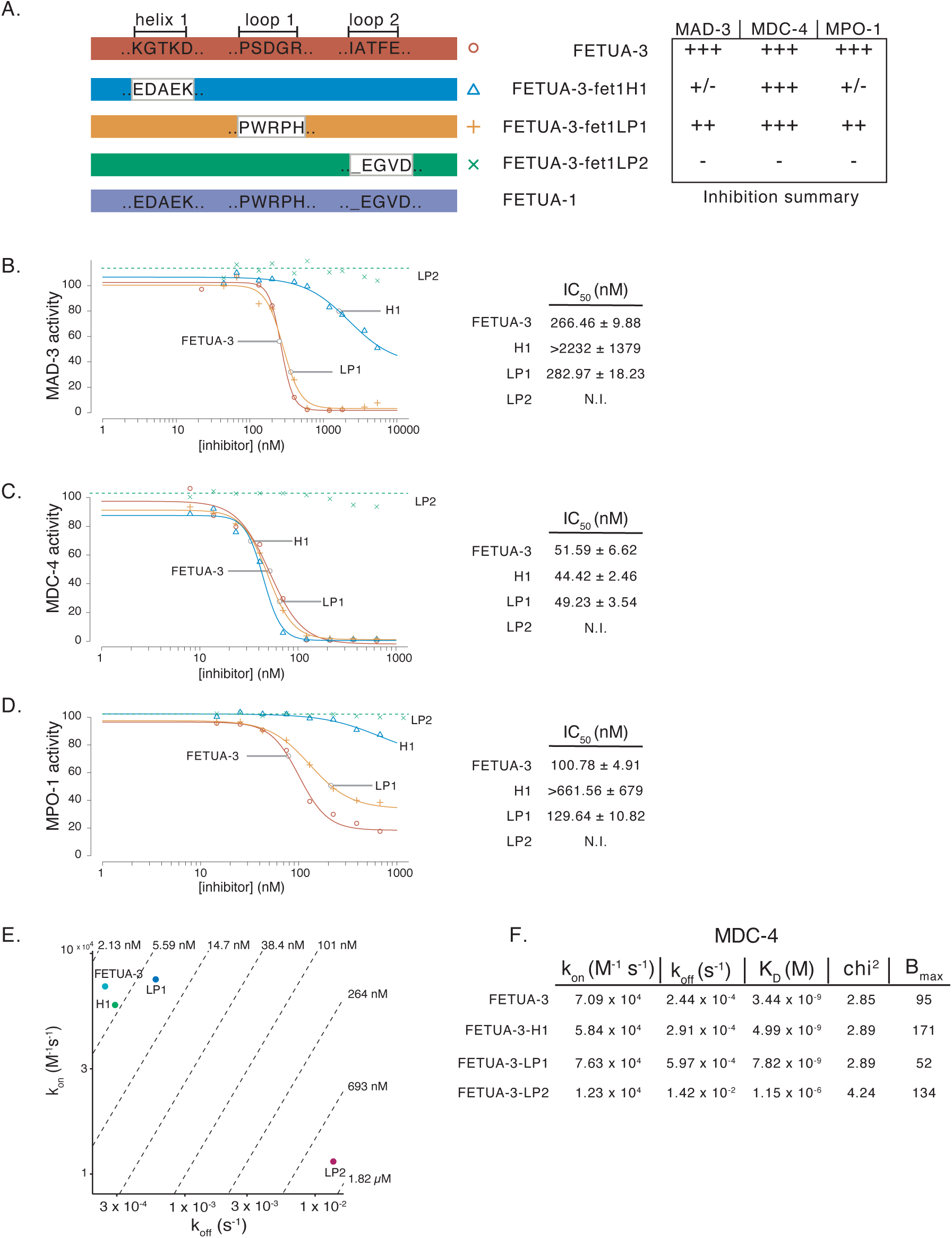
The H1 and Loop2 FETUA domains are required for multi-target inhibition. (A) A schematic representation of the chimeric FETUA-3 proteins tested for inhibition of and binding to MAD-3, MDC-4 and MPO-1. The FETUA-3 amino acid sequences for helix 1 (H1), loop 1 (LP1) and loop 2 (LP2) are shown in black letters in the top rectangle (red). Those sequences were individually swapped for the respective FETUA-1 sequences shown in the bottom rectangle (purple). The chimeric FETUA-3-fet1H1 (blue rectangle) consists of the FETUA-3 protein with the H1 sequence replaced with FETUA-1 amino acids. The chimeric FETUA-3-fet1LP1 (orange rectangle) and FETUA-3-fet1LP2 (green rectangle) replace the FETUA-3 LP1 and LP2 sequences with the respective FETUA-1 LP1 amino acids. An indel at the start of LP2 also shortens LP2 in FETUA-3-fet1LP2 relative to the length of FETUA-3 LP2. A summary of the inhibition data presented in B – D is shown to the left of the protein schematics and uses the protein names to label the rows of the table. (B-D) Inhibition of MAD-3 (B), MDC-4 (C) and MPO-1 (D) by FETUA-3, FETUA-3-fet1H1 (H1), FETUA-3-fet1LP1 (LP1) and FETUA-3-fet1LP2 (LP2). The colors of the lines are matched to the rectangles in (A) and the midpoints (IC_50_ ± standard error, nM) of the curves are reported in the tables on the right. N.I. = no inhibition is detected. (B) Replacement of LP1 sequences slightly increases inhibition of MAD-3 by FETUA-3, whereas replacement of H1 strongly reduces (IC_50_ > 2232 nM) and replacement of LP2 abolishes inhibition. (C) Replacement of LP1 or H1 sequences has no or slight effects on FETUA-3 inhibition of MDC-4, but replacement of LP2 abolishes inhibition. (D) Replacement of LP1 modestly reduces, while replacement of H1 or LP2 abolishes inhibition of MPO-1 by FETUA-3 (E) An iso-affinity plot shows the ∼334-fold reduction of the FETUA-3-fet1LP2 -MDC-4 equilibrium dissociation constant (K_D_ = 1.15 x 10^-6^ M) is due to a slower on-rate (k_on_) and a faster off-rate (k_off_) relative to FETUA-3 (K_D_ = 3.44 x 10^-9^ M). (F) A table showing association and dissociation rate constants and equilibrium dissociation constants for MDC-4 interactions with FETUA-3 and FETUA-3-fet1H1, FETUA-3-fet1LP1 and FETUA3-fet1LP2.

We found that FETUA-3-fet1H1 does not inhibit MAD-3 (Figure 4B, IC_50_ > 2232 nM) or MPO-1 (Figure 4D, IC_50_ > 662 nM) but efficiently inhibits MDC-4 (Figure 4C, IC_50_ ∼ 44 nM). We also find that FETUA-3-fet1H1 binding to the respective MPs is consistent with its inhibitory activity. The MDC-4::FETUA-3-fet1H1 and MDC-4::FETUA-3 binding constants are similar (Figure 4F, K_D_: FETUA-3-fetH1= 4.99 x 10^-9^ M, FETUA-3 = 3.44 x 10^-9^ M and Figure S6A) but binding between FETUA-3-fet1H1 and MAD-3 or MPO-1 is severely reduced (K_D_ > 10^-6^ M; Figures S6B, C). This data highlights key differences in how FETUA-3 interacts with different MPs such that some modifications (the C9G substitution and the HI swap) that minimally disrupt binding to or inhibition of MDC-4 severely reduce or eliminate binding to and inhibition of MAD-3 or MPO-1.

In contrast to these strong, differential effects of modifying helix1, we observed only slight effects upon replacing loop 1 on FETUA-3 inhibition of and binding to all three enzymes. The chimeric protein FETUA-3-fet1LP1 showed similar activity to FETUA-3 in its ability to inhibit MDC-4 (Figure 4C), a slight 1.06-fold decrease in activity in inhibiting MAD-3 (Figure 4B) and a small 1.3-fold decrease in the inhibition of MPO-1 (Figure 4D). We also observed decreases in FETUA-3-fet1LP1 binding to all three enzymes relative to FETUA-3 including a 2.3-fold effect on binding to MDC-4 (K_D_ = 7.82 x 10^-9^ M vs. K_D_ = 3.4 x10^-9^ M; Figure 4E and F) and ∼7-fold effects on binding to MAD-3 (K_D_ = 101 x 10^-9^ M) and MPO-1 (K_D_ = 176 x 10^-9^ M) FETUA-3 (Figure S7A-C). We conclude that loop 1 contributes a small degree to target binding when other FETUA-3 domains are intact.

However, we found that replacement of FETUA-3 “loop 2” (FETUA-3-fet1LP2) abolished inhibition of all three enzymes (Figures 4B-D), eliminated the detection of binding to MAD-3 and MPO-1 (Figure S8B, C), and severely reduced binding to MDC-4 (Figures 4E, F and Figure S8A). The complete loss of inhibition activity and consistent effect on binding to all three enzymes suggests that “loop 2” is a core element of FETUA-3 that is necessary for multi-target inhibition. The location of “loop 2” in the predicted structure model is distant from the enzyme active site so one possible explanation for these results is that “loop 2” is primarily involved in the binding of FETUA-3 to multiple MPs. We do detect some binding of FETUA-3-fet1LP2 to MDC-4 (K_D_ = 1.15 x 10^-6^ M), however this is a ∼350-fold reduction in affinity relative to wild-type FETUA-3 (K_D_ = 3.44 x 10^-9^ M; Figure 4E, F; Figure S7G, H). The reduction in the binding constant is largely due to a ∼57-fold increase in the off-rate suggesting a role for “loop 2” in stabilization of the inhibitor-complex.

These results demonstrate that two additional FETUA-3 structural elements, the H1 region and loop 2, are required for multi-target binding and inhibition, and thus these two interfaces also changed during the evolution of FETUA-3 from the ancestral FETUA-1 protein. To further elucidate the mechanism of enzyme inhibition, we next examined the possible roles of individual NTD residues.

### The first residue of the N-terminal domain is critical for multi-target inhibition

The AlphaFold models predict that the phenylalanine residue at position one of the FETUA-3 NTD is very near the catalytic site of MAD-3b and raises the possibility that the residue could influence the electrostatic and metal-coordination environment of the Zn²⁺-containing active site. We also note that this residue is strictly conserved in all FETUA-3 sequences throughout the 50-million-year radiation of vipers (Figure 5A)(30). Therefore, we hypothesized that the hydrophobic benzyl side chain could also contribute to binding through hydrophobic interactions and that its bulky, planar structure could sterically hinder access to the active site.

**Figure 5.**
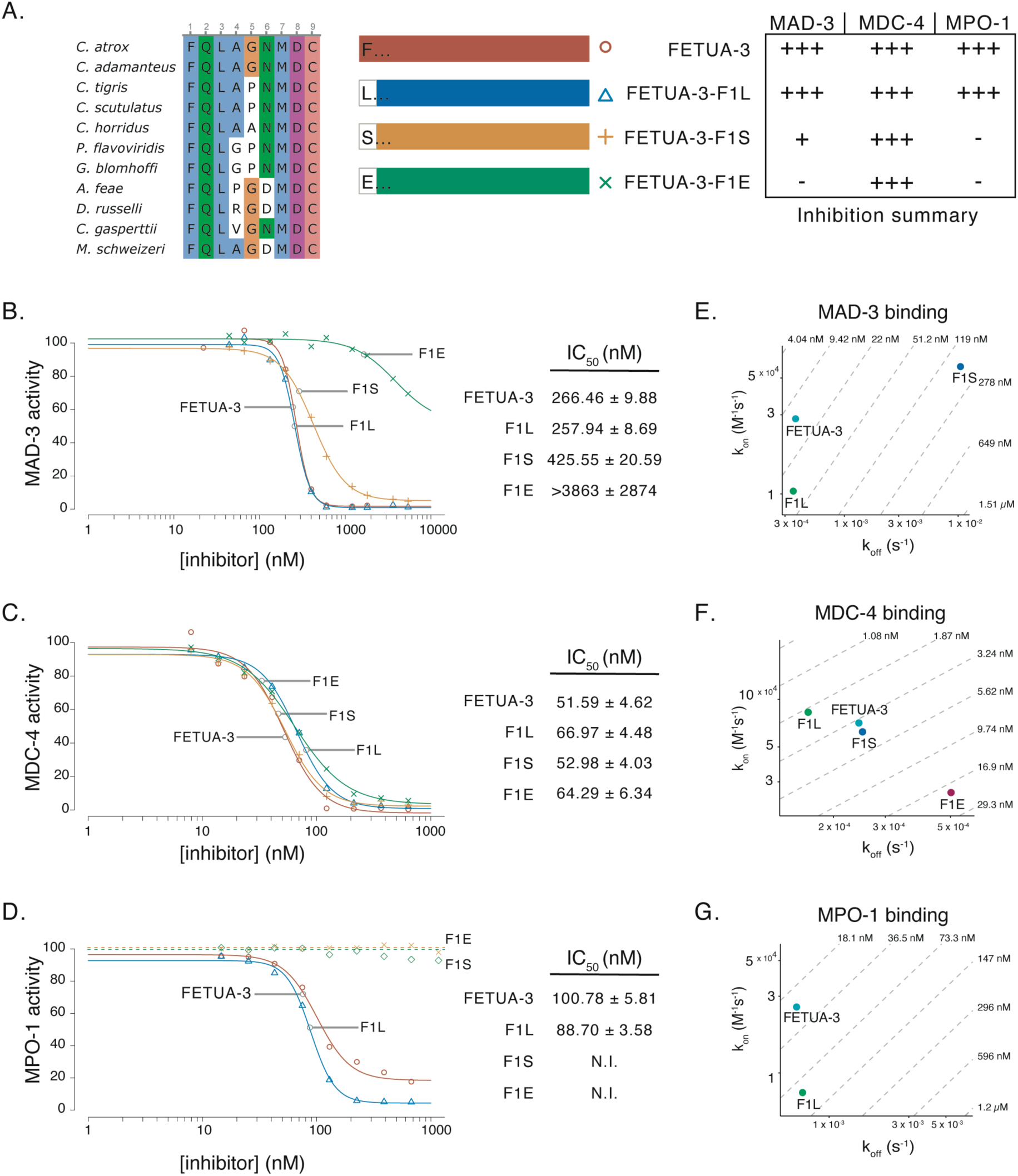
The first residue of the N-terminal domain is critical for multi-target inhibition. (A) Left, an alignment of FETUA-3 NTD protein sequences from pit viper and true vipers species which shared a common ancestor ∼ 50 million years ago shows strict conservation of the phenylalanine-1, glutamine-2 and leucine-3 residues. Right, a schematic representation of the FETUA-3 proteins containing a single amino acid substitution at position one (F1) that were tested for inhibition of and binding to MAD-3, MDC-4, and MPO-1. A summary of the inhibition data presented in B – D is shown to the left of the protein schematics and uses the protein names to label the rows of the table. (B, C, D) Inhibition of MAD-3 (B), MDC-4 (C), and MPO-1 (D) by FETUA-3 (red line), F1L (blue), F1S (orange) and F1E (green) with the midpoints (IC_50_ ± standard error, nM) of the curves reported in the tables to the right. The F1L substitution has minimal or undetectable effects on FETUA-3 inhibition of all three enzymes. (B) The F1S substitution has a slight effect on FETUA-3 inhibition of MAD-3, but the F1E substitution strongly reduces inhibitory activity (IC_50_ > 3863 nM). (C) The F1S and F1E substitutions have no or very slight effects on FETUA-3 inhibition of MDC-4 activity. (D) The F1S and F1E substitutions abolish inhibitory activity of MPO-1. (E, F, G) Binding to MAD-3 (E), MDC-4 (F) and MPO-1 (G) by FETUA-3 and F1 mutants. (E) The FIE substitution reduces binding of FETUA-3 to greater than 1 µM (see methods regarding limits for binding measures in this study). The FETUA-3-F1S binding constant to MAD-3 is reduced ∼19-fold. (F) The FIE substitution reduces binding of FETUA-3 to MDC-4 by ∼6-fold. (G) The F1S and FIE substitutions eliminate FETUA-3 binding to MPO-1 (K_D_ > 1µM), and thus accounting for the loss of MPO-1 inhibition.

To test the contribution of the hydrophobic side chain while removing the aromatic character of Phe, we replaced Phe1 (F1) with leucine (FETUA-3-F1L; Figure 5A). We observed only very slight effects on the inhibition of (Figures 5B-D), or binding to all three MPs by FETUA-3-F1L (Figures 5E-G).

The modest effects of the F1L substitution prompted us to replace the nonpolar phenylalanine with a polar serine residue thus substituting the benzyl ring with a polar hydroxyl group (Figure 5A, F1S). Relative to wild-type FETUA-3, FETUA-3-F1S inhibited MAD-3 (IC_50_: WT = 266, F1S = 426 nM) and MDC-4 (IC_50_: WT = 52, F1S = 53 nM) with slightly reduced or similar potency (Figure 5B, C). However, inhibition of MPO-1 activity was abolished (Figure 5D, orange dashed line).

To further dissect the effect of the F1S substitution we compared the interaction kinetics of FETUA-3-F1S with the three MPs. We observed a small (1.2-fold) decrease in binding to MDC-4 (K_D_: WT = 3.44 x 10^-9^ M, F1S = 4.03 x 10^-9^ M), a larger (19-fold) decrease in binding to MAD-3 (K_D_: WT = 1.3 x 10^-8^ M, F1S = 24.5 x 10^-8^ M) and no binding to MPO-1 (Figures 5F, E, G and Figure S9A - C). The decrease in binding affinity between FETUA-3-F1S and MAD-3 can be attributed mainly to a faster off-rate (k_off_ : WT = 3.72 x 10^-4^ s^-1^, F1S = 1.0 x 10^-2^ s^-1^) suggesting that the presence of serine at position one of the N-terminus is permissive for initial recognition of MAD-3 by FETUA-3 (Figure 5E, Figure S9B). The undetectable binding of FETUA-F1S to MPO-1 indicates that FETUA-3 binding to and inhibition of MPO-1 operates under stricter constraints.

The discovery that a change of the first residue of the NTD from phenylalanine to serine has drastic effects on MPO-1 inhibition and binding lead us to explore whether a more radical substitution of phenylalanine with glutamic acid could disrupt inhibition of the other MPs (Figure 5A). The F1E substitution abolished FETUA-3 inhibition of and binding to MAD-3 (IC_50_ > 3863 nM) as well as MPO-1 (Figures 5B, D; Figure S10A - C). Surprisingly, FETUA-3-F1E inhibited MDC-4 nearly as well as wild-type FETUA-3 (IC_50_: WT = 52, F1E = 64 nM; Figure 5C) while binding to the enzyme was decreased five-fold (K_D_: WT = 3.44 x 10^-9^ M, F1E = 19.5 x 10^-9^ M; Figure 5F, Figure S10A).

The effects of the substitutions of phenylalanine at position one of the NTD reveal several facets of FETUA-3 function. First, the selective loss of MPO-1 inhibition by the FETUA-3-F1S protein, and of MPO-1 and MAD-3 inhibition by the FETUA-3-F1E protein demonstrates there are differences among MP target enzymes that likely impose constraints on the side chain that is permissible at position one of the NTD, and thus may account for the strict evolutionary conservation of Phe1. Second, all effects on MP inhibition are accounted for by effects on MP binding, that is, no side chain substitution uncouples target binding from inhibition. And third, all three different side chains are inhibitory with respect to MDC-4 activity, suggesting a limited role for the benzyl side-chain ring in enzyme inhibition. To gain further insight into the mechanism of MP inhibition we revisited what is known about snake venom metalloproteinases’ mechanism of action and the AF3 models of FETUA-3-MP interactions.

### Backbone elements of Phe1 disrupt catalysis

It has been proposed that catalysis by snake venom metalloproteases involves a Zn²⁺-bound water molecule that is oriented and activated by a conserved catalytic Glu residue(40). This positioning of the water molecule enables nucleophilic attack on the substrate carbonyl group and subsequent cleavage of the peptide bond. In Michaelis complexes, the substrate carbonyl oxygen is also in close proximity to the Zn²⁺, positioning the scissile bond for nucleophilic attack(40).

Detailed inspection of the MAD-3b active site in the AF3 models of its complex with FETUA-3 reveals that the primary amine of the Phe1 residue of FETUA-3 is located in close proximity (∼2.9Å) to the carboxylate group of the catalytic Glu, indicating a potential salt bridge. In addition, the carbonyl oxygen of Phe1 is ∼1.8Å from the Zn^2+^ ion (Figure 6A and Figure S13A), a distance compatible with zinc coordination. Together, these interactions raise the possibility that Phe1 may influence the active-site environment of MAD-3b, potentially altering the availability of a water molecule involved in catalysis.

**Figure 6.**
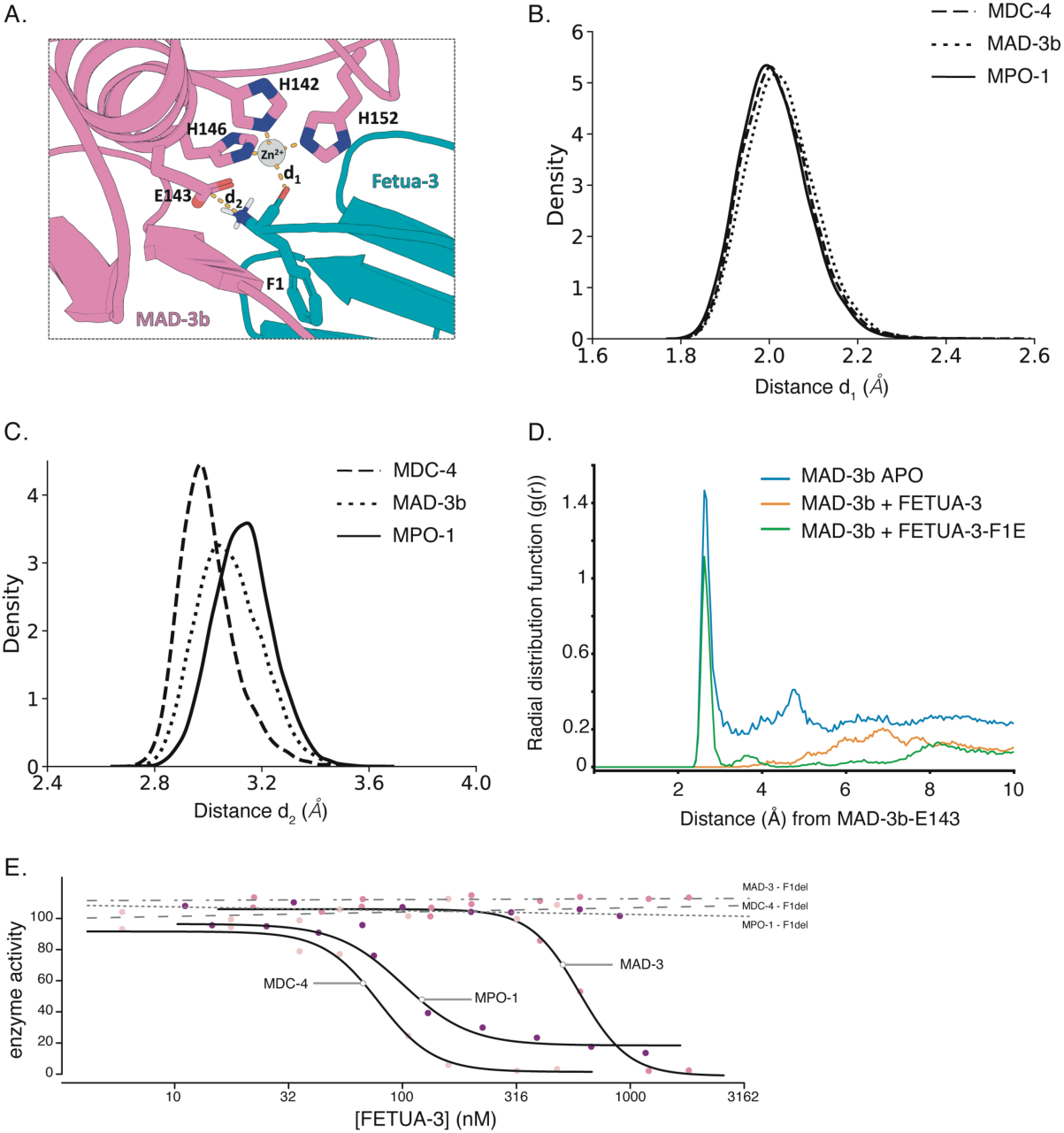
Two backbone elements of Phe1 are critical for MP inhibition. (A) The MAD-3b (pink) active site showing side chains of the catalytic residues: glutamic acid (E143) and three histidines 142, 146 and 152 (H142, H146, H152) are shown coordinating a zinc ion (grey sphere). Two core elements of the FETUA-3-F1 residue (blue-green) likely contribute to MP inhibition: the carbonyl oxygen (red) coordination of the zinc ion and the alpha-amine (blue) forming a salt-bridge with E143 of the MP. The distances between the core elements are designated d1 (carbonyl oxygen to zinc) and d2 (amine group nitrogen to carboxylate carbon of E143). (B-C) Probability distributions of the distances from the QM/MM MD simulations between two core elements of the N-terminal Phe1: (B) the carbonyl oxygen and Zn^2+^ (d1) and (C) the amine nitrogen and carboxylate carbon of the catalytic Glu (E143) (d2). Distances for MDC-4, MAD-3b and MPO-1 are shown with dashed, dotted and full lines, respectively. (D) Radial distribution function (RDF) from three MM MD simulations showing the presence of water (peak ∼2.6 Å) in the MAD-3b APO MD (blue) and MAD-3b::FETUA-3-F1E MD (green) and no water density detected in the MAD-3b::FETUA-3 (orange). (E) Deletion of Phe1 abolishes FETUA-3 inhibition of MDC-4 (dashed line), MAD-3 (dot-dashed line), MPO-1 (dotted line). FETUA-3 (solid lines) inhibition curves for of MDC-4, MAD-3 and MPO-1 are shown for reference.

With respect to Phe1 and the potential mechanism of inhibition of MDC-4 and MPO-1, we could not discern mechanistic insights from the original, low confidence AF3 MDC-4 and MPO-1 complex models. In these models the global orientation of FETUA-3 relative to the enzymes resembles that in the MAD-3b complex (Figure 2A, left and right panels), however, the binding interfaces differ substantially from the MAD-3b complex. In both predicted complexes key molecular features (NTD and loop2) adopt poses incompatible with inhibition, with Phe1 and the NTD positioned far from the active site and loops 1 and 2 arranged substantially farther from the enzyme than in the MAD-3b model (Figure S11A,B). Because our experimental results demonstrate that both the NTD and loop2 of FETUA-3 are required for binding to and inhibition of MDC-4 and MPO-1, we reject the original MDC-4 and MPO-1models.

We then explored whether the predicted and experimentally supported MAD-3b::FETUA-3 complex could serve as a template for modeling the other two MP::FETUA-3 complexes. We superimposed MDC-4 and MPO-1 onto MAD-3b and transferred the FETUA-3 coordinates from the MAD-3b::FETUA-3 complex (see Materials and Methods). We then assessed the original and refined models with physics-based metrics that are independent of the criteria used to build AF models(41) by scoring each complex with Rosetta InterfaceAnalyzer(42) across five independent relaxation replicates.

The refined models recover 70 to 96% of the difference from the MAD-3b reference in interface completeness and shape complementarity, and replicate variation falls by more than half (Figure S12). The resulting interfaces are thus physically reasonable, internally reproducible, and consistent with the functional data. In both of the refined MDC-4 and MPO-1 complexes with FETUA-3, the primary amine of Phe1 is near the carboxylate group of the catalytic Glu, and its carbonyl oxygen is near the Zn^2+^ ion, as in the MAD-3b complex (Figure S11).

The AF3 models provide a static view of FETUA-3 within the active site. To further investigate how Phe1 might disrupt metalloproteinase cleavage, we turned to Quantum Mechanics/Molecular Mechanics (QM/MM) molecular dynamics (MD) simulations on each complex of MDC-4, MAD-3b, and MPO-1 with FETUA-3 in the inhibitory pose (see Methods). Three independent replicas, 1 ns each, were run at the DFTB3-3OB/MM level of theory(43). All MD trajectories remained stable throughout the simulation time, with no significant drift observed in backbone RMSD, supporting the reliability of the sampled conformational ensembles (Figure S13B). The analysis of the MD simulations revealed three key features of the complex: i) coordination of the Zn^2+^ by the backbone carbonyl oxygen of the first residue of FETUA-3 (Phe); ii) salt-bridge formation between the catalytic glutamic acid (Glu) and the primary amine of Phe-1; and iii) the absence of water within the catalytic site. These interactions persist throughout the QM/MM MD simulations and thus provide support for the proposed binding mode and indicate they remain dynamically stable over the course of the simulations.

With respect to coordination of the Zn^2+^ ion, we observe distances around 2Å between Zn^2+^ and the epsilon nitrogens (Nε) of the three metalloproteinase histidines across all simulated systems. Additionally, the probability distributions of the distances between the carbonyl oxygen (=O) of FETUA3-F1 and Zn^2+^ are similarly centered around ∼2 Å for all three MPs (Figure 6B), supporting a model with a coordination of the catalytic zinc by the N-terminal carbonyl oxygen. We also observe that the average distance between the FETUA3-F1 amino group (NH₃⁺, which we infer to be protonated based upon a pKa of ∼8.0) and the carboxylate (COO^-^) of the metalloproteinase glutamic acid to be ∼3.0 Å (Figure 6C), consistent with a strong electrostatic salt-bridge interaction.

To further investigate the observation of the absence of water within the catalytic site of the enzyme when bound by FETUA-3, we also ran classical MM MD simulations over a longer time scale. We used the equilibrated portion of MM MD simulations (Figure S13C) to derive the RDF of water around catalytic Glu143 (Figure 6D). The radial distribution function (RDF) shows a clear density of water molecules at ∼2.6 Å from the catalytic Glu143 in the apo MAD-3b, consistent with the catalytic water proposed in the reaction mechanism(40). In contrast, no comparable water density was observed near Glu143 in the MAD-3b–FETUA-3 complex over the same timescale consistent with the inhibition (Figure 6D). To explore whether the absence of water could instead reflect the more restricted access and limited simulation timescale relative to the apo enzyme, we also simulated the F1E mutant in complex with MAD-3b. Interestingly, the F1E complex showed a clear water density near Glu143 comparable to that observed in apo MAD-3b on the same timescale (Figure 6D), consistent with the experimentally observed loss of inhibition by FETUA-3-F1E (see Figure 5B).

In summary, the MD simulations support a model of inhibition in which FETUA3 Phe1, with a bulky hydrophobic side chain, sterically restricts solvent access to the active site, coordinates Zn^2+^ via the backbone carbonyl oxygen, and promotes a strong salt bridge between the N-terminal amine and the carboxylate of the catalytic glutamate, thus preventing substrate cleavage.

If this model of the interactions of the Phe1 residue in the MP catalytic site is accurate, then the removal of this residue might affect FETUA-3 activity in different ways than the replacement of Phe1 with amino acids bearing different side chains. Indeed, we observe that a mutant FETUA-3 protein in which Phe1 is deleted (FETUA-3-F1del) completely lacks inhibitory activity against all three MP enzymes, including MDC-4 (Figure 6D). To determine whether the loss of activity was independent of or also entailed loss of FETUA-3 binding to MPs we also measured the binding kinetics of FETUA-3-F1del to each MP enzyme. We found that FETUA-3-F1del binding to MDC-4 was severely reduced (∼465-fold reduction; K_D_ = 16.7 x 10^-6^ M) and binding to MAD-3 and MPO-1 was undetectable (K_D_ > 10^-6^ M) (Figure 6E and Figure S14A - C). These results reveal a critical role for Phe1 in FETUA-3 function and are consistent with the Phe side chain being required for multi-target binding and the backbone elements of the residue directly contributing to inhibition of catalysis by all three enzymes.

### Multi-target inhibition imposes functional and sequence constraints on the FETUA-3 N-terminus

In view of the essential role of Phe1 and its strict evolutionary conservation, we also investigated the contributions of the two strictly conserved, adjacent residues glutamine-2 (Gln2, Q2) and leucine-3 (Leu3, L3) to FETUA-3 function.

AlphaFold modeling suggests that the amide group of the Gln2 side chain may hydrogen bond with residues in target MPs (for example, Ile-108 of MAD-3b; Figure S15A), so we interrogated the potential requirement for Gln2 by making an alanine substitution (FETUA-3-Q2A; Figure 7A) and measuring its effects on enzyme inhibition and binding. The inhibition of MAD-3 and MDC-4 by FETUA-3-Q2A was comparable to FETUA-3 (Figures 7B, C). However, MPO-1 was only weakly inhibited by FETUA-3-Q2A (Figure 7D, IC_50_ > 3150 nM). In binding assays, the FETUA-3-Q2A on-rates were similar to that of FETUA-3 (Figure7E, F, G; Figure S16A - C) but the off-rates were 6.9-, 1.9-, and 9.4-fold faster for MAD-3, MDC-4 and MPO-1, respectively (Figures 7E, F, G; Figure S16A - C). The faster dissociation of FETUA-3-Q2A from all three MPs suggests that glutamine in the second position may be necessary to stabilize the inhibitor-enzyme complex for all MP targets. However, it is puzzling that there are differences in the extent of the loss of inhibition (severe for MPO-1 and minimal for MAD-3) given their similar off-rates.

**Figure 7.**
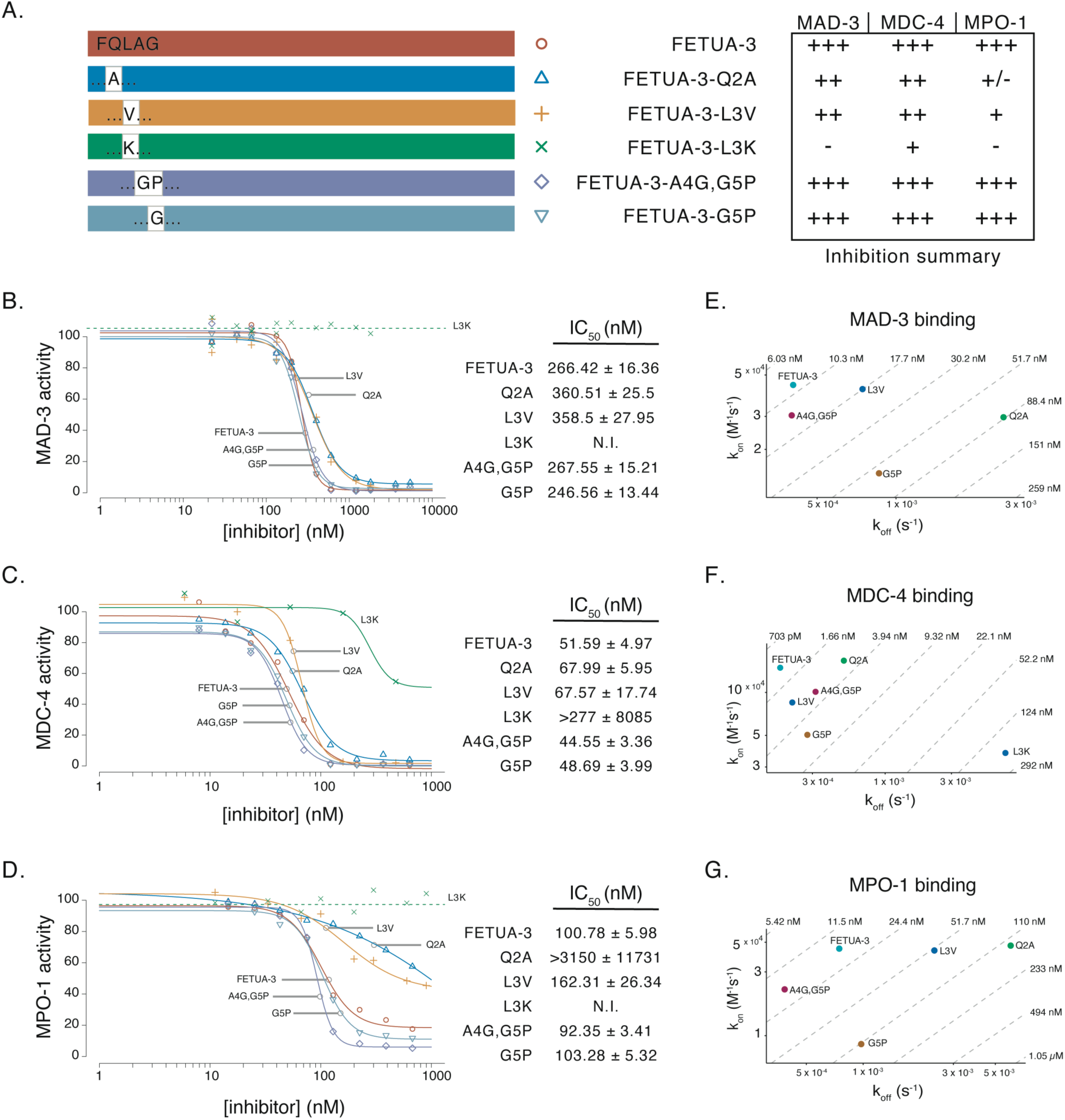
Functional constraints restrict the sequence composition of the N-terminal domain. (A) A schematic representation of the FETUA-3 proteins containing single amino acid substitutions at positions two through five that were tested for inhibition of and binding to MDC-4, MAD-3 and MPO-1. A summary of the inhibition data presented in B – D is shown to the left of the protein schematics and uses the protein names to label the rows of the table. (B, C, D) Inhibition of MAD-3 (B), MDC-4 (C), and MPO-1 (D) by FETUA-3 (red line), FETUA-3 -Q2A (blue), FETUA-3-L3V (orange), FETUA-3-L3K (green), FETUA-3-A4G,G5P (purple) and FETUA-3-G5P (light blue) with the midpoints (IC_50_ ± standard error, nM) of the curves reported in the tables to the right. (B) FETUA-3-L3K does not inhibit MAD-3 whereas the Q2A, L3V, A4G/G5P, and G5P substitutions have slight or no effect on enzyme inhibition. (C) FETUA-3-L3K weakly inhibits MDC-4, whereas the Q2A, L3V, A4G/G5P, and G5P substitutions have slight or no effect on enzyme inhibition. (D) FETUA-3-L3K does not inhibit MPO1, and the Q2A and L3V substitutions also have strong effects on MPO-1 inhibition. The A4G/G5P and G5P substitutions have slight effects on MPO-1 inhibition. (E - G) Binding of MAD-3 (E), MDC-4 (F) and MPO-1 (G) by FETUA-3 (blue-green point), and the Q2A (green), L3V (blue), L3K (blue), A4G, G5P (magenta) and G5P (brown) mutant proteins. (E) FETUA-3-L3K binding to MAD-3 is not detected. (F) The L3K substitution reduces FETUA-3 binding to MDC-4 ∼17.6 fold (K_D_ = 20.8 x 10^-9^M) whereas binding by the other mutant FETUA-3 proteins is only slightly affected. (G) FETUA-3-L3K binding to MPO-1 is not detected, while FETUA-3-Q2A and FETUA-3 L3V binding to MPO-1 are reduced, with a more pronounced effect observed on the off-rates (faster).

Analysis of predicted structures suggested that Leu3 may also contact residues in target MPs (Ile-170 of MAD-3b) in a central position of a hydrophobic “wedge” at the FETUA-3::MP interface (Figure S15B). To investigate the role of L3 in FETUA-3 function we made proteins containing a conservative valine substitution (FETUA-3-L3V) or a non-conservative lysine substitution (FETUA-3-L3K) and tested their activities (Figure 7A and S17A - C). We find that FETUA-3-L3V inhibits MAD-3 and MDC-4 activity similarly to wild-type FETUA-3 (Figures 7B, C) but observe weak and incomplete inhibition of MPO-1 (Figure 7D, IC_50_ > 162 nM). This contrasts with FETUA-3-L3K which does not inhibit either MAD-3 and MPO-1 and only weakly inhibits MDC-4 (IC_50_ > 277 nM; Figures 7B, D, C). The effects of the L3K substitution on MP inhibition appear to be explained by the effects on MP binding as binding to MAD-3 and MPO-1 are abolished (Figure S18A - C) while binding to MDC-4 is reduced ∼57-fold (L3K: K_D_ = 195 x 10^-9^ M). Kinetic measurements of binding to target MPs by FETUA-3-L3V reveal minimal perturbation of association rates (k_on_) but an increasingly faster dissociation rate for MDC-4, MAD-3 and MPO-1, respectively (Figures 7E, F, G; rightward shift of L3V). This observation suggests that the third residue of the NTD likely functions in stabilizing the FETUA3::MP inhibitory complex more so than initial target binding.

In contrast to the strict evolutionary conservation of the first three residues of the FETUA-3 N-terminus, there is sequence divergence among vipers at amino acid positions 4 and 5. For example, Ala4 in *C. atrox* FETUA-3 is a glycine residue in several pit viper species and Gly5 is a proline residue in some pit viper species, and both residues are substituted in several species (Figure 5A). To test the effects of these naturally-occurring amino acid substitutions on FETUA-3 activity, we made recombinant forms of *C. atrox* FETUA-3 in which Gly5 was replaced with proline (FETUA-3-G5P) and Ala4 and Gly5 were replaced with glycine and proline, respectively (FETUA-3-A4G, G5P). We found that both FETUA-3-G5P and FETUA-3-A4G,G5P inhibit all three MP targets as well or nearly as well as FETUA-3 (Figures 7B, C, D) and bind to each target with very similar kinetics (Figures 7E, F, G). Therefore, the strict evolutionary conservation of N-terminal residues 1-3 of FETUA-3 reflects very strict functional constraints required to maintain multi-target inhibition while the interspecific variation at positions 4 and 5 reflects more relaxed functional constraints on these sites.

## Discussion

The radiation of vipers was advanced by the expansion and diversification of venom metalloproteinases, which was accompanied by the coevolution of the FETUA family of MP inhibitors, a key biochemical innovation that enabled the venomous lifestyle. In this study, we have identified several structural features of the rattlesnake FETUA-3 protein that are required for the inhibition of a diverse set of venom metalloproteinases. These findings enable us to address three main questions of interest. First, how did FETUA-3 evolve from a non-inhibitory ancestral Fetuin-A into a potent, multi-target MP inhibitor? Second, to what extent is FETUA-3’s mechanism of action novel with respect to other types of vertebrate MP and protease inhibitors? And third, how does a small number of FETUA proteins control the activity of a large set of venom toxins? The answers to these questions bear on our general understanding of how new high-affinity protein-protein interactions and protein networks arise.

### The molecular evolution of FETUA-3 function entailed the modification of three protein interfaces

FETUA-3 evolved from the ancestral FETUA-1 (Fetuin-A) glycoprotein during the early evolution of vipers through the process of gene duplication and divergence(30). Since FETUA-1/Fetuin-A is not an inhibitor of venom metalloproteinases, or of any known metalloproteinase or protease, FETUA-3’s function constitutes a biochemical novelty. While there is a substantial body of work on how proteins evolve novel specificities for small molecule ligands, substrates, or sites within larger molecules (7,44,45), there are relatively few instances in which we understand how the more extensive and complex interfaces evolve between proteins. Therefore, it is of general interest to identify the scope of molecular changes involved in the *de novo* evolution of FETUA-3 function (i.e. did the multi-target MP binding and inhibitory functions require one, a few, or many mutations?).

Structure prediction indicated that there are three major FETUA-3-MP interfaces with each having several putative pairwise residues in contact. Our experimental results established that these three structural elements of the FETUA-3 protein are required for MP binding and inhibition. Importantly, our rational mutagenesis approach included the swapping of conserved regions of FETUA-1 with the homologous, conserved regions of FETUA-3, as well as the substitution of individual residues in the NTD. Therefore, we are able to infer from these experiments that the acquisition of FETUA-3 function involved numerous substitutions across each of three regions, comprising:

1. The N-terminal domain, consisting of eight residues adjacent to the first cysteine (C9, conserved in all FETUA proteins), that inserts into MP active sites. The FETUA-3 NTD sequence FQLAGMND is strongly conserved among vipers (Figure 5) and differs at most (6 of 8) positions from the well-conserved FETUA-1 NTD sequence HHLQSQID.
2. A segment of the first alpha helix (H1) consisting of residues 12-16, replacement of which was found to disrupt MP regulation in the order of MPO-1>MAD-3> MDC-4 (strongest to weakest effect). The FETUA-3 H1 sequence KGTKD is perfectly conserved among vipers and differs at all sites from the EDAEK sequence that is conserved among FETUA-1 proteins.
3. Loop 2, consisting of residues 99-103, and shown to be required for FETUA-3 binding to and inhibition of all three MPs. Three residues within the *C. atrox* FETUA-3 sequence **I**A**TF**E (shown in bold) are well-conserved among vipers and differ at all positions from the _EGVD sequence that is conserved in most FETUA-1 proteins.

The number of residues within these three essential regions that differ from FETUA-1 and were subsequently strongly conserved in FETUA-3 totals fourteen, thirteen of which are predicted to make contacts with two or all three target MPs. This is a much larger number of contacts and functional substitutions than those involved in major shifts in substrate or ligand specificity (Siddiq et al. 2017). We suggest that this larger number of substitutions reflects the longer evolutionary path and a greater degree of molecular tuning required for the *de novo* evolution of high affinity binding to and inhibition of multiple MP proteins.

This inference is consistent with a few studies that have explored the evolution of new protein subunit interactions(46), the evolution of high affinity protein complexes (47), and the affinity maturation of antibody-protein complexes (48). It is also consistent with what is known about protein-protein interactions in general. The predicted FETUA-3::MP interfaces involve a range of 37-47 contact residues encompassing an interface area of 1700-2100 Å^2^. A classic survey of many categories of protein complexes found that the typical protein-protein interaction involves about 50 residues and an overall interface area of about 1600 (± 400) Å^2^ (8). Thus, the *de novo* evolution of high affinity protein-protein interfaces generally requires changes to a large number of residues.

### The mechanism of FETUA-3 action is novel with respect to other vertebrate MP inhibitors

Since FETUA-3 is a novel MP inhibitor, it is also of particular interest to understand to what degree FETUAs evolved a new solution or exploited a pre-existing solution for inhibiting MPs. That is, which of FETUA-3’s functional features are shared with or distinct from mechanisms employed by the two other known classes of inhibitors of the several large families of vertebrate MPs or the many types of protease inhibitors?(49)

Taken together, structure prediction and experimental data suggest that the FETUA-3 mechanism of action entails: i) target binding to regions adjacent to active sites via Loop2 (HP2) and H1; ii) target binding within active sites via electrostatic interactions through the insertion of the N-terminus into the active site; iii) positioning of the N-terminus in an inhibitory pose by the C9-C296 disulfide bond and iv) disruption of catalysis via interactions of the Phe1 primary amine with the MP catalytic Glu residue (Glu 143 or Glu 144) and coordination of Zn^2+^ by the carbonyl oxygen of Phe1 (Figure 8A; N for primary amine, O for carbonyl oxygen, NTD for N-terminal domain, H1 for helix1 and HP2 for Loop2).

**Figure 8.**
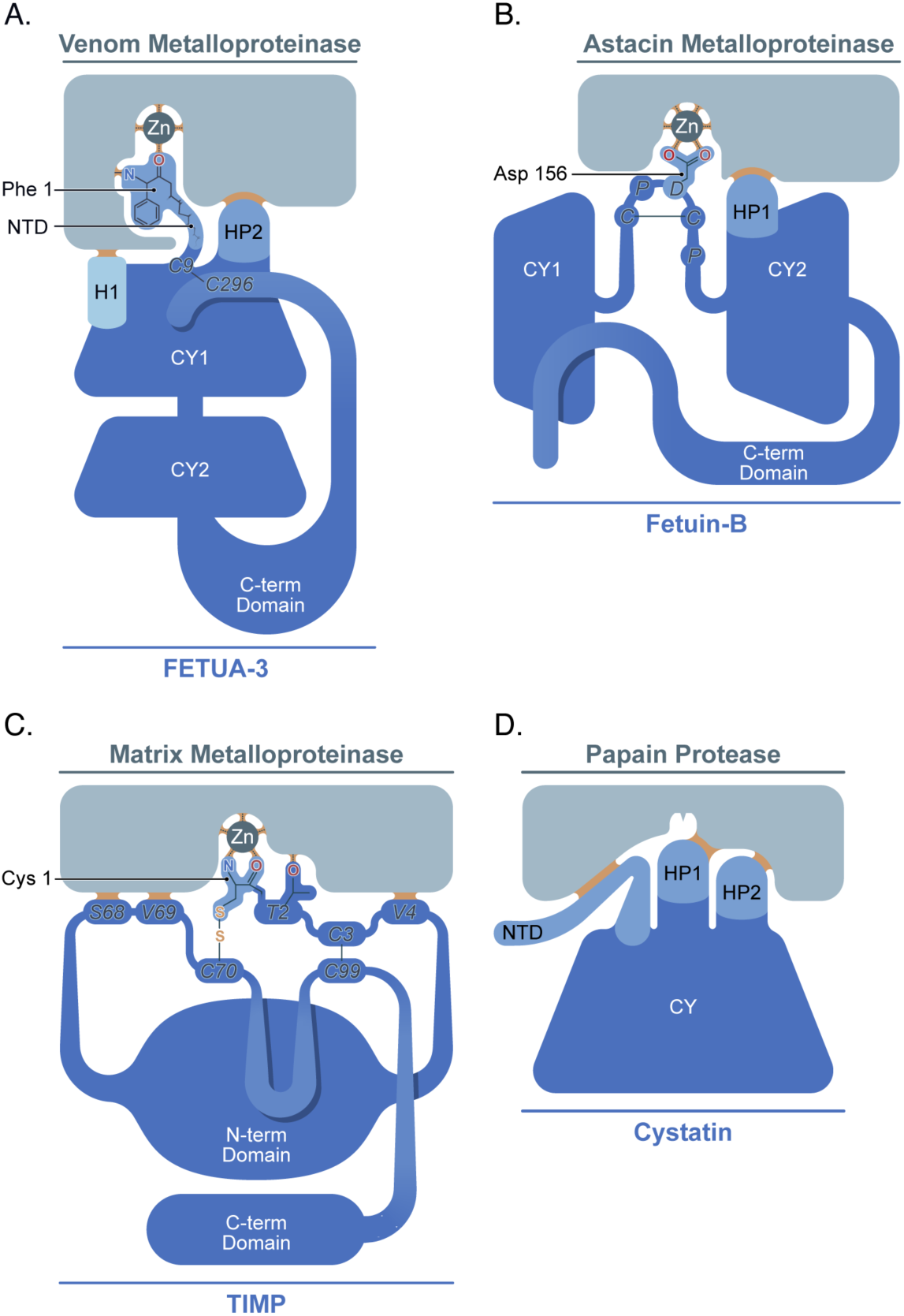
FETUA-3 convergently evolved a unique mechanism of MP inhibition. (A-D) Schematic diagrams highlighting the mechanisms of protease binding and inhibition by (A) FETUA-3; (B) Fetuin-B; (C) TIMPs; and (D) cystatins. The respective molecules, domains and active sites are not drawn to scale. (A) FETUA-3 binds to venom MPs through its N-terminal domain (NTD), and helix 1 (H1) and second hairloop (HP2) domains within its first cystatin domain (CY1). Backbone elements, nitrogen (N) from N-terminal amine and oxygen (O) from carbonyl group, of the Phe1 residue inhibit catalysis via interactions with the catalytic glutamate and Zn^2+^ within target MP active sites. The disulfide bridge (C9-C296) likely stabilizes the NTD in a binding and inhibitory posture. (B) Despite a similar domain organization, Fetuin-B binds to targets via different cystatin domains (CY2) and inhibits the catalytic machinery via different structural elements (CPDCP linker) than FETUA-3. (C) TIMPs utilize backbone elements of their N-terminal residue (cysteine-1, C1) to inhibit the catalytic machinery (a glutamate residue and zinc) in a manner similar to FETUA-3. The N-terminus is also stabilized by a disulfide bond to a distal cysteine (C70). However, TIMPs bind to their respective targets through entirely different interfaces (V4, S68, V69) and are not evolutionarily related to FETUA-3. (D) Cystatins bind to target proteases (but not zinc metalloproteinases) through hairpin loops (HP1 and HP2) and insert a portion of their N-terminal domain (NTD) into target active sites, however, they do not affect the catalytic machinery.

While we identify some analogies between FETUA-3’s mechanism of action and those of Fetuin-B(50) (an astacin and meprin MP inhibitor), TIMPs(26) (MMP and ADAM inhibitors), or cystatins(51) (cysteine protease inhibitors), we conclude that FETUA-3 represents a distinct class of MP inhibitors. For example, at the protein domain level, FETUA-3 possesses two tandem cystatin domains, a feature consisting of an alpha helix lying on top of an anti-parallel beta sheet. This domain also occurs in tandem in FETUIN-B, a paralog of FETUIN-A. However, while the key H1 and Loop2 motifs occur in the first cystatin domain of FETUA-3, in Fetuin-B the first cystatin domain contributes little to binding while the second cystatin domain and a linker region between the tandem cystatin domains are crucial for target binding and inhibition(50) (Figure 8B). Moreover, the mechanisms of enzyme inhibition by these two inhibitors are completely distinct. Our results indicate that the backbone elements (alpha-amine and carbonyl oxygen) of the FETUA-3 N-terminal Phe1 directly disrupt the catalytic machinery. In contrast, it is the side chain of an aspartate residue (Asp 156) in the “raised-elephant-trunk” formed by the linker region of Fetuin-B that binds to the catalytic zinc of target enzymes(50) (Figure 8B).

The FETUA-3 mechanism of disrupting catalysis exhibits some resemblance to that of TIMPs in that these MP inhibitors also employ an N-terminal residue (Cys1 in the case of TIMPs) whose primary amine interacts with a catalytic Glu residue and backbone carbonyl oxygen group binds to the catalytic zinc and displaces the critical water molecule(52–57). The N-terminal residue is positioned in the catalytic site by a disulfide bond with a distal cysteine (Cys 70; Figure 8C)(58). These molecular analogies are instructive in terms of revealing a common strategy of effectively inhibiting zinc metalloproteinases (binding to the catalytic glutamate and coordination of the catalytic zinc), however, they were attained through entirely independent evolutionary paths and different structural determinants. Therefore, FETUA-3 represents a case of convergent functional evolution.

Finally, we note that cystatins also bind to target proteases through two hairpin loops and insert a portion of their N-terminal domain into target active sites, however they do not affect the catalytic machinery(49,51) (Figure 8D).

### Multi-target inhibition: Master keys for different locks

The most important functional properties of individual FETUA proteins, with respect to both their biological roles in snakes and potential therapeutic utility, is their inhibition of multiple and diverse venom MPs. The MP toxin family has diversified rapidly among vipers such that even different species of rattlesnakes possess widely varying numbers of MP genes (5-30 genes;(22,59–62) and express different combinations of individual venom MPs belonging to 15-16 paralog groups(22,62) whose metalloproteinase domain sequences diverge considerably (∼30-50%). Yet, rattlesnakes and other vipers have co-evolved just 4-5 FETUA paralogs (30). This ratio of a small number of FETUA proteins to a large number of diverse MPs indicates that individual FETUAs likely regulate numerous MPs, which has been demonstrated for C. atrox FETUA-3 (29) and FETUA-2 from certain Asian pit vipers (63). Thus, understanding the structural requirements for multi-target inhibition is important for understanding FETUA function and FETUA-MP co-evolution.

We have shown here that various substitutions in the strictly conserved N-terminal sequences of FETUA-3 cause the loss of inhibition of selective MP targets. In particular, various substitutions that cause the loss of inhibition of MAD-3 and/or MPO-1 have slight or no effect on inhibition of MDC-4. Since the FETUA-3 N-terminal domain inserts into the active site, the differential effects of the N-terminal amino acid substitutions on target binding and inhibition reveals that there are differences among MP active sites that must be accommodated by FETUA-3.

If we use a lock and key analogy for the inhibitor and target enzymes, FETUA-3 is a master key that fits slightly different locks. The very strict conservation of the three N-terminal amino acid residues thus reflects the strict structural constraints necessary to fit a subset of MPs with related but slightly different locks (active sites) and maintain multi-target inhibition by FETUA-3.

It is important to underscore that FETUA-3 does not fit all locks, there are other master keys for other sets of locks. For example, FETUA-3 does not regulate rattlesnake MPs primarily responsible for causing hemorrhage, but FETUA-2 and FETUA-5 do(30). Conversely, FETUA-2 does not regulate MDC-4, MAD-3, or MPO-1(29). In addition, different combinations of FETUAs are required to fully neutralize the lethal effects of different species’ venoms, which further indicates that different FETUA proteins act on different subsets of MPs(30). All rattlesnake and viper FETUA paralogs have a similar domain organization as FETUA-3, but their N-terminal domain sequences are distinct from one another and highly conserved(30). Based on the findings here, we anticipate that the different FETUA N-terminal domains may be essential to their respective MP target specificities. Therefore, it will be valuable to examine other FETUA-MP interactions in order to elucidate the molecular basis of FETUA-MP specificity and to determine whether all FETUAs inhibit MPs through a similar mechanism.

### From structure prediction to mechanism

In light of the growing capabilities of and interest in structure prediction methods, particularly as a potentially more expeditious approach to understanding protein-protein interactions than physical structure determination(63,64), we thought it might be useful to comment briefly on the powers and limitations of these methods in our experience here. With respect to powers, multiple methods yielded high confidence structures for FETUA-3-MAD-3b and revealed several interfaces that were then experimentally demonstrated to be essential for FETUA-3-MP binding and inhibition. On the other hand, these methods failed to predict high confidence structures for FETUA-3::MDC-4 and FETUA-3::MPO1 complexes, despite these being high affinity interactions that were shown experimentally to be dependent on the same interfaces as MAD-3.

This experience reveals that it was fortunate that we included three biochemically validated FETUA-3 targets in this study. Had MAD-3b not been available for study, we would not have been able to identify candidate functional features of FETUA-3. Similarly, had the failures of the MDC-4 and MPO-1 predictions deterred us, we would not have discovered the common features necessary for FETUA-3 inhibition of all three MPs.

The reasons for the wide variation in confidence among different predicted FETUA-3-MP structures are not certain. One possibility is that the venom MPs are challenging, and in fact such disagreement is characteristic of the snake venom MP protein family. A systematic comparison across more than 1,000 snake venom toxins lacking experimental templates found that snake venom MPs displayed the largest variability between modeling approaches of any toxin family, with discrepancies of up to 145 Å between methods and agreement largely restricted to the conserved peptidase domain(65). The same study found that current methods perform well for structured domains but remain unreliable for flexible regions such as loops and propeptides. Another possibility is that although the FETUA-3 structure is fairly well predicted, it is a novel MP inhibitor and there is no precedent in training datasets for its mode of interaction. The wide variation in prediction scores with established FETUA-3-MP targets suggest that caution is warranted in exploring previously uncharacterized protein-protein interactions solely with *in silico* targets.

## Materials and Methods

### Recombinant FETUA protein production

FETUA-3 and FETUA-1 wild-type and recombinant constructs and proteins were prepared commercially (GenScript, Piscataway, NJ, USA). The sequences encoding *C. atrox* FETUA-1, FETUA-3 and its variants proteins were based on sequences obtained from the genome of one *C. atrox* specimen(30). To facilitate their expression as recombinant proteins, the endogenous signal peptide sequence (1-19 amino acids) was replaced with a sequence optimized for protein expression (MGWSCIILFLVATATGVHS) in mammalian cell lines. The synthetic genes were cloned into the mammalian expression vector pcDNA3.4 (Invitrogen, Carlsbad, CA, USA).

The recombinant protein expression plasmids were transiently transfected into CHO-S cells and grown in serum-free media to facilitate the isolation of the tag-free proteins. After five days, the 40-100 ml culture supernatants were harvested and the proteins (isoelectric points approximately 5.6-5.8) were isolated by applying the supernatants to a HiTrap Q Fast Flow anion-exchange column (Cytiva) that was equilibrated with a low salt loading buffer (20mM Tris pH 8.0, 0% NaCl (w/v)) and eluting with a linear salt gradient (20mM Tris pH 8.0, 0-30% NaCl (w/v). The purity of the recombinant proteins was analyzed by size-exclusion chromatography on a TSKgelG3000SWXL HPLC column developed with 0.1M Na_2_SO_4_ in 0.118 M phosphate buffer pH 7.0 and estimated to be 95-99%. The samples were dialyzed against either Tris-buffered saline (TBS) or phosphate-buffered saline (PBS) and stored at -80°C.

### Monoclonal antibody generation

Mouse monoclonal antibodies targeting FETUA-3 were generated commercially (Genscript) by immunization of mice with using the same GST-FETUA-3 fusion antigen (containing residues 78-268) previously described(29). A panel of ten monoclonal antibodies reactive with FETUA-3 were then screened for cross-reactivity with FETUA-1, FETUA-2, FETUA-4, and FETUA-5 by enzyme-linked immunosorbent assay (ELISA). One antibody that cross-reacted with all five FETUA proteins (16E12) was selected for recombinant expression and production by a commercial vendor (Genscript). Briefly, the antibody V_H_ and V_L_ domain regions were cloned and sequenced, then fused to a IgG2a constant region, expressed in Chinese Hamster Ovary cells, and purified from the culture supernatant by affinity chromatography on Protein A. The antibodies were then tested for their utility in Surface Plasmon Resonance experiments described below.

### Structural modeling of FETUA-3–metalloprotease complexes

For structure prediction, truncated MPs comprising the first 202 residues were used. MP::FETUA-3::Zn²⁺ complexes were predicted independently using AlphaFold 3 (AF3), OpenDDE and ESMFold2, with all predictions performed locally on an HPC cluster. For AF3, model parameters were obtained upon request from Google DeepMind. OpenDDE predictions were generated with the general-purpose base checkpoint (opendde.pt), intended for non-antibody biomolecular complexes. In all methods, the MP and FETUA3 sequences, as well as Zn²⁺, specified as a ligand using the corresponding CCD code, were provided as JSON input. For each method and each complex, 25 independent seeds were used, generating five structural models per seed (125 predictions per complex), with 10 recycling iterations. All other parameters, including the number of diffusion steps, were kept as default. All predicted structures, together with their associated output metrics, are provided in the corresponding Zenodo directory [10.5281/zenodo.21921413].

### Rosetta Refinement and Interface Analysis

The five highest-ranked AF3 structures for each MP::FETUA3::Zn²⁺ complex were subsequently refined using the Rosetta FastRelax protocol (Rosetta v3.12). The Rosetta-refined structures were subsequently analyzed using the Rosetta InterfaceAnalyzer protocol to characterize the MP–FETUA-3 interfaces. We report the predicted binding energy (dG_separated, in Rosetta Energy Units, REU); the total solvent-accessible surface area buried upon complex formation (dSASA_int, Å²); the binding energy normalized by buried interface area (dG_separated/dSASAx100, REU per 100 Å²); shape complementarity (sc_value, ranging 0–1); the number of cross-interface hydrogen bonds (hbonds_int); the change in the number of buried unsatisfied hydrogen-bond donors and acceptors upon complex formation (delta_unsatHbonds); and the number of interface residues (nres_int). Both FastRelax and InterfaceAnalyzer were performed using the ref2015 scoring function. The interface analysis was performed between chains A and B excluding zinc from the interface analysis.

### Caseinolytic Assay

Protease activity was measured using Pierce Protease Assay Kit (Thermo Fisher Scientific), according to manufacturer’s instructions with slight modifications as described below. Purified enzymes were used at the following concentrations calculated based on predicted molecular weights: MDC-4 (500ng, 53nM), MAD-3 (1 µg; 200.8 nM), and MPO-1 (500 ng; 100nM). Pre-incubation (at 37°C) of enzymes with molar excess FETUA-3 was allowed to reach equilibrium before adding casein (150 µg for MDC-4 and MPO-1, 200 µg for MAD-3) and allowing substrate cleavage to proceed for one hour. The reaction was developed by addition of 50 µl of trinitrobenzenesulfonic acid (0.03 % TNBSA) and incubated at room temperature for forty minutes. Absorbance was measured at 450 nm on a microplate reader. The assays were carried out in triplicate. In the initial characterization of FETUA-3 inhibition we noted the half-maximal inhibitor concentration (IC_50_) approached the concentration of the respective enzyme which suggested our experimental conditions may reveal that FETUA-3 is a tight binding inhibitor. To further test if FETUA-3 is a tight binding inhibitor, we repeated the FETUA-3 inhibition titrations while varying the enzyme concentration and observed an increase in the IC_50_ with increasing enzyme concentration suggesting FETUA-3 is a tight binding inhibitor(66). Thus, we fit the FETUA-3 inhibition data to the Morrison equation using GraphPad to obtain inhibition constants (K_i_)(67). For the relative comparison of FETUA-3 enzymatic inhibition to FETUA-3 mutants, data was fit to the four-parameter logistic model and the inhibitor concentration at which half of the enzymatic activity is inhibited was identified using the drc R-package(68). The low affinity for casein by MAD3 (highest K_M_) limited the practicality of substantially increasing substrate (casein) concentration to monitor the effect on IC_50_ therefore the experiment was only performed with MDC-4 and MPO-1.

### Surface Plasmon Resonance binding analysis

The binding kinetics between MDC-4, MAD-3 and MPO-1 and FETUA-3 recombinant proteins were measured using the Alto digital SPR system (Nicoya). Preliminary results showed the FETUA-3 and FETUA-1 were stably retained during SPR experiments therefore a FETUA-3 capture surface was generated with the anti-FETUA monoclonal antibody 16E12. The antibody (20 μg/mL in 10mM Sodium acetate, pH 4.1) was immobilized on a 16-channel carboxyl cartridge (Nicoya KC-CBX-CMD-16) using the amine-coupling method (EDC/NHS). FETUA-3 (ligand) capture was performed by exposing the FETUA recombinant proteins to the immobilized antibody surface (300 seconds, expected RU of 100 - 200). Single cycle kinetics (SCK) of the analyte (MDC-4, MAD-3, MPO-1) - ligand interaction was measured by applying five concentrations, in a 3-fold dilution series, of the analyte (MDC4: 0.74 nM – 60 nM) over at least two ligand concentrations (FETUA-3: 100 nM and 200 nM). For each enzyme a minimum of two broad concentration ranges to characterize interactions between “strong” (nanomolar concentrations) and “weak” (micromolar concentrations) binders. Interactions were measured in TBST-Ca (TBS pH7.4 containing 0.1% Tween-20 and 2mM CaCl_2_) at 37°C. Association and dissociation times were 180 and 600 seconds, respectively. Sensor surfaces were regenerated between SCK runs using 10mM glycine-HCl, pH 2.5.

SPR responses to each analyte were calculated by subtracting the response from a reference sensor channel (not exposed to ligand). Sensograms were fitted to a 1:1 Langmuir binding model using TraceDrawer software (https://tracedrawer.com). For FETUA-3 and MDC-4, MAD-3 and MPO-1 interactions kinetics parameters were obtained from three data sets with chi² value less than 10% of the corresponding Rmax (Bmax). Data was visualized using ggplot2(69) and ggrecipes(70).

### Binding analysis and data presentation

Undetected binding could be due to perturbation of specific sequences that are necessary for binding or because the range of analyte tested was below the specific limit for detecting weak interactions. To exclude the latter option a “reference protein” was included in kinetic experiments that aimed to detect weak binding. This reference serves as an internal experimental control and shows the weakest binding interaction reproducibly detected at the highest analyte concentrations (MDC-4: 0.12 – 10 µM, MAD-3: 0.56 – 15 µM, MPO-1: 0.25 – 20 µM). The binding constant derived from the reference protein serves as an upper limit of detection for weak binders. Additionally, to verify all FETUA proteins are immobilized (regardless of binding activity) at relatively equivalent levels the response units (RU) after antibody capture are shown as individual points (unique color for each protein). These plots of immobilization levels and binding curves with reference proteins are shown in supplementary figures.

### MM MD Simulations

All MM MD simulations were performed on the previously described AlphaFold-predicted models using Amber24 using pmemd.cuda engine(71). For the molecular mechanics (MM) MD simulations, the Amber ff19SB force field(72) was employed and water molecules were modeled using the OPC model(73).

The Zn²⁺ coordination sphere was reparameterized following the standard non-bonded model in the Amber24 tutorial using the Python-based Metal Center Parameter Builder (MCPB.py)(74). Briefly, the three coordinating histidines, Zn²⁺ and a water molecule were optimized at the B3LYP/6-31G* level of theory. In the optimized structure, the electrostatic potential (ESP) was calculated at the same level of theory. Atomic partial charges for the metal coordination sphere were then fitted to this electrostatic potential. During the fitting procedure, the charge of the coordinated water molecule was constrained to zero, allowing it to be later replaced by the inhibitor without introducing inconsistencies in the charge distribution. The resulting mol2 files are available in the Zenodo repository (10.5281/zenodo.21921413).

All of the systems were prepared using tleap in Amber24(71). Disulfide bridges in the MPs and FETUA-3, suggested by AF3 predictions, were manually defined as bonds in tleap. Protonation states of titratable residues were assigned using PropKa at pH 7.5(75). Na⁺ and Cl⁻ ions were added to neutralize the system and achieve a physiological ionic strength of 150 mM. The simulation box was solvated so that the distance from the protein-inhibitor complex to the box edge was 12 Å.

Each system was first minimized using 1000 steps of steepest descent (SD) followed by 200 steps of conjugate gradient (CG) minimization. Systems were then gradually heated from 100 to 310 K over 1 ns using a Langevin thermostat (collision frequency γ = 2 ps⁻¹) in the NPT ensemble, with pressure controlled by a Monte Carlo barostat. During heating, a positional restraint of 100 kcal·mol⁻¹·Å⁻² was applied to the protein.

After heating, systems were equilibrated for 1 ns, while reducing the restrain weight on the protein from 100 to 10 kcal·mol⁻¹·Å⁻². A further minimization of 1000 steps using SD and 30 steps of CG was performed with restraints only on protein backbone atoms, allowing the side chains to adjust. The system was then equilibrated in the NPT ensemble, while gradually reducing restraint weights by 2.5 kcal·mol⁻¹·Å⁻² for 1 ns. A final 1 ns of unrestrained equilibration was performed prior to production runs.

Production MD simulations were carried out in the NVT ensemble using a 2 fs timestep using the SHAKE algorithm(76) to constrain bonds involving hydrogen. Electrostatic interactions were computed using particle-mesh Ewald(77),while a 10 Å cutoff radius was applied for all nonelectrostatic interactions. Temperature was maintained at 310 K using the Langevin thermostat (γ = 2 ps⁻¹). Three independent replicas of 250 ns each were run starting from different initial velocity seeds.

### QM/MM MD Simulations

QM/MM MD simulations were performed starting from the equilibrated MM MD structure using the Amber24 pmemd.MPI engine(71). In all the simulated systems, the QM region consisted of the three coordinating histidines (cut between C_α_-C_β_), the catalytic Zn²⁺ and the N-terminal residue of FETUA-3 (Phe1 in the wild-type, or the corresponding residue in each mutant). The QM region included the entire N-terminal residue of FETUA-3, with the QM/MM boundary placed at the second residue (between the Cα and carbonyl carbon and between Cα and Cβ).

The QM region was described using the DFTB3/3OB semiempirical functional, which is suitable for Zn²⁺ coordination sites(43,78). The remainder of the protein and solvent were described using the ff19SB force field(79) and OPC water model(73), respectively. To validate the choice of QM method and avoid potential artifacts, additional simulations were performed using GFN2-xTB(80).

The cutoff for both long-range MM interactions and QM/MM electrostatic interactions was set to 15 Å. The system temperature was maintained at 310 K using a Langevin thermostat (collision frequency γ = 2 ps⁻¹). Three independent replicas were run for each system, starting from different random velocity seeds.

### MD Trajectory Analysis

Root-mean-square deviations (RMSD), root-mean-square fluctuations (RMSF),interatomic distances, angles and radial distribution functions (RDFs) were calculated using cpptraj in Amber24(71). All plots were generated using custom Python scripts.

## Data Availability

The Molecular Mechanics and Molecular Dynamics data files are available in the Zenodo repository (10.5281/zenodo.21921413)

## Acknowledgments

We thank Adamo Mancino of the HHMI/Janelia cryo-EM facility for his efforts to obtain structures of protein complexes, Robert Rutherford of Nicoya Lifesciences for his help with SPR experiments, Kate Baldwin for help with illustrations, and Matt Giorgianni, Yetunde Ayinuola, and Jory van Thiel for comments on the manuscript. We thank Dr. Steffen Schmidt for his support with the local compute infrastructure at the Department of Biochemistry, University of Bayreuth, and the HPC FAU Erlangen for support and resources within the ProtDynDesign project. M.A. would like to thank Prof. Iñaki Tuñón (University of Valencia) for fruitful discussions on MD simulations and analysis. The R.F.L. laboratory is supported through funding from the National Institute of Health Carlos III to CNIO. S.V.T. was supported by the “Generación D” initiative for talent attraction (C005/24-ED CV1, Red.es, Ministerio para la Transformación Digital y de la Función Pública) funded by the European Union NextGenerationEU funds, through PRTR. Modeling and computational analysis was supported by institutional funds of the University of Bayreuth (B.H.). This work was supported by the Howard Hughes Medical Institute (S.B.C.) and the Andrew and Mary Balo and Nicholas and Susan Simon Endowed Chair at the University of Maryland (S.B.C).

## Author Contributions

N.L.D., F.P.U, M.H., and E.C. conducted biochemical experiments; S.V.T. and R.F.L. conducted structure prediction analyses; M.A. and B.H. conducted structural modeling of complexes, QM/MM and MD runs, and Rosetta refinement and analyses; N.L.D., S.V.T., M.A., and S.B.C wrote the manuscript in consultation with all authors.

## Ethics Declaration

Competing Interest Statement: S.B.C. and F.P.U are co-inventors on patents pending from related work, of which the University of Maryland is the assignee (U.S. Patent App. No. 63/431,147 and 63/994,480).

