## Supplemental Information for "The *de novo* molecular evolution of a viper venom metalloproteinase inhibitor and its target network"

### The novel viper FETUA-3 protein evolved a unique mode of inhibiting snake venom metalloproteinases

‡ Co-first authors

*Corresponding author: Sean B Carroll.

Figure S1 A - C

A. MDC4 digestion of succinylated casein

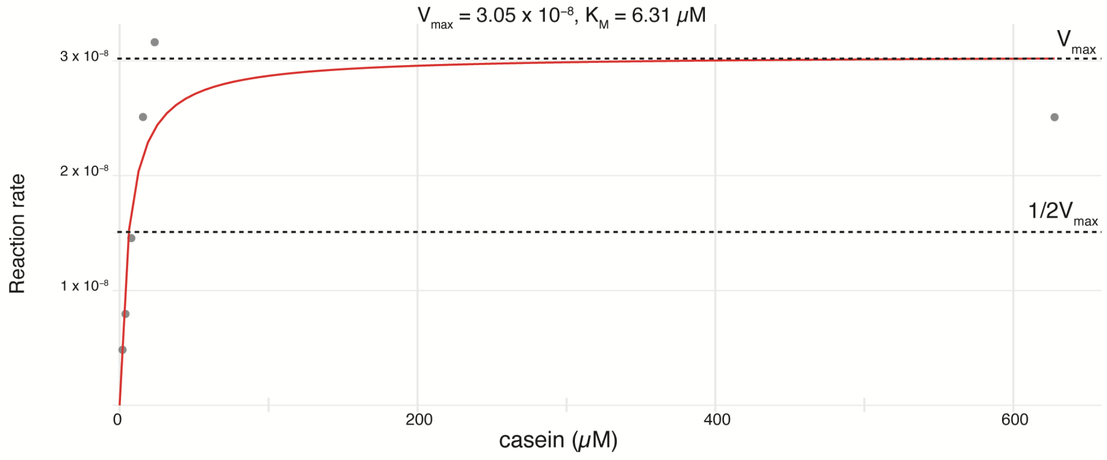


B. MAD3 digestion of succinylated casein

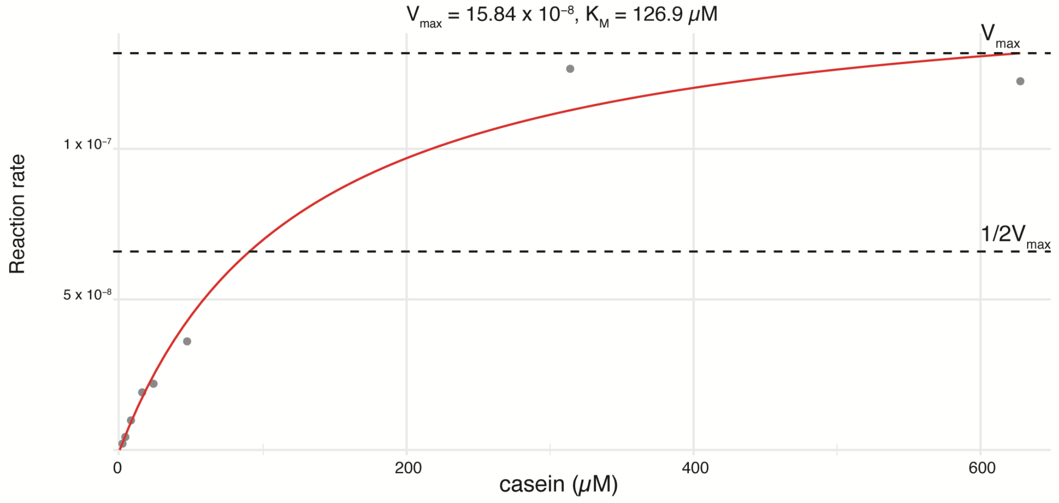


C. MPO1 digestion of succinylated casein

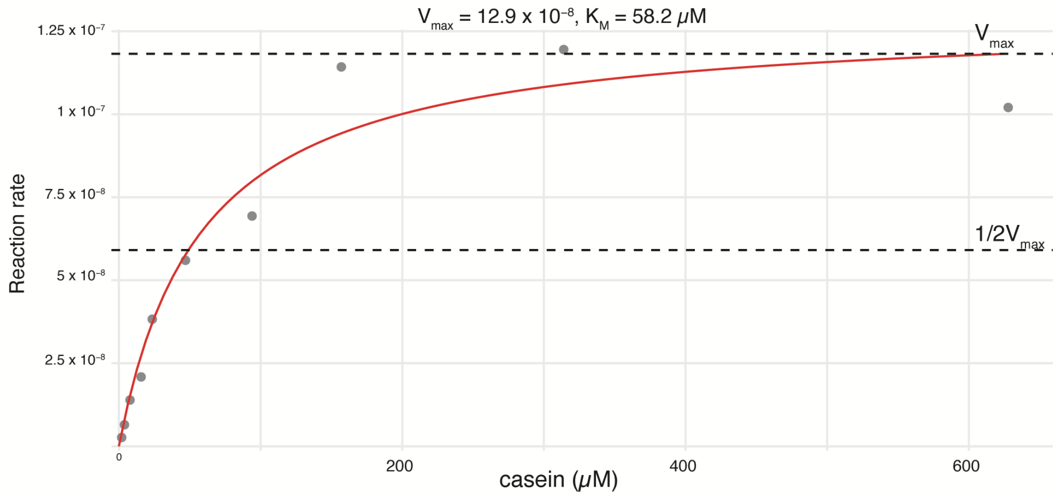

**Figure S1. Venom metalloproteinase digestion of succinylated casein.**

For MDC-4 (A), MAD-3 (B) and MPO-1 (C) the initial rate (velocity, v) of substrate cleavage was determined for eight concentrations (1.96 to 628 μM) of casein. Substrate concentration versus reaction rate was fit to the two parameter Michaelis-Menton model.

Figure S2 Distributions of ipTM and pTM metrics for predictions of FETUA-3 with MAD-3b, MPO-1 or MDC-4 across three methods.

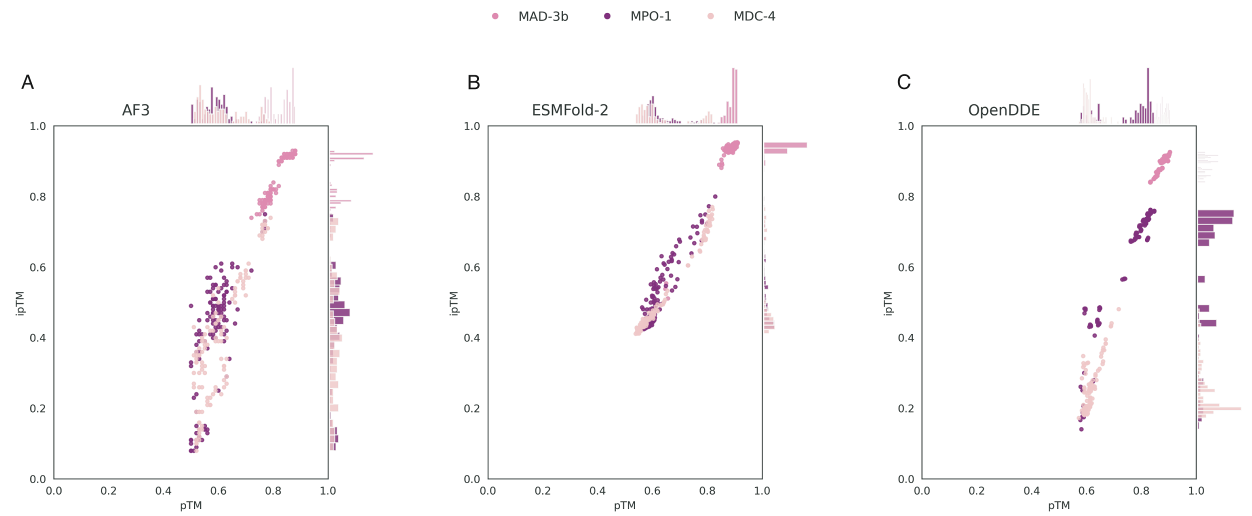


| | ipTM | pTM | ipTM | pTM | ipTM | pTM |
|---|---|---|---|---|---|---|
| MAD-3b | 0.86 ± 0.06 | 0.82 ± 0.04 | 0.92 ± 0.09 | 0.88 ± 0.05 | 0.90 ± 0.03 | 0.87 ± 0.02 |
| MDC-4 | 0.40 ± 0.18 | 0.61 ± 0.08 | 0.52 ± 0.09 | 0.63 ± 0.09 | 0.25 ± 0.07 | 0.61 ± 0.02 |
| MPO-1 | 0.42 ± 0.16 | 0.59 ± 0.06 | 0.52 ± 0.09 | 0.63 ± 0.06 | 0.60 ± 0.19 | 0.73 ± 0.10 |

**Figure S2. Distributions of ipTM and pTM values for models of FETUA-3 with MAD-3b, MDC-4 and MPO-1 generated using three structure prediction methods.**

We generated 125 models per complex from 25 independent seeds, each producing five diffusion samples using AlphaFold-3, ESMFold-2 and OpenDDE. (A) AlphaFold-3 predictions of FETUA-3 with MAD-3b clustered together (dark pink points, top right corner) and yielded high confidence scores (average ± standard deviation of ipTM 0.86 ± 0.06, pTM 0.82 ± 0.04). Confidence scores of FETUA-3 complexes with MDC-4 (light pink points) and MPO-1 (magenta points) were spread across a wide range of low confidence. Average ipTM and pTM scores are shown below each plot. In general, FETUA-3::MAD-3b complexes were predicted with high confidence with all three methods while the FETUA-3::MDC4 and FETUA-3::MPO1 complexes were predicted with relatively lower confidence.

Figure S3 Agreement between and within three structure prediction models

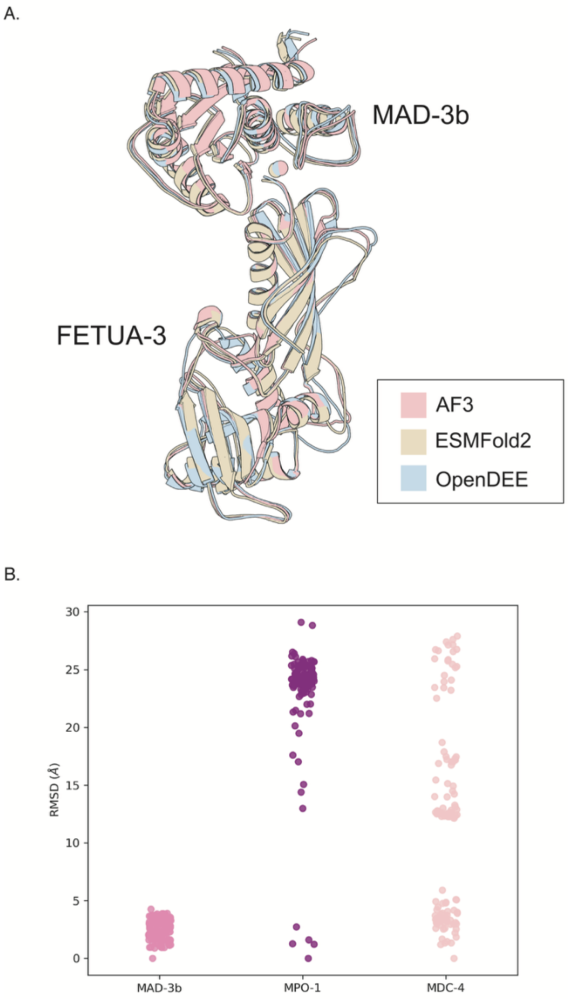


**Figure S3. Agreement between and within three structure prediction methods for the FETUA-3–MAD-3b complex**.

(A) The OpenDDE and ESMFold2 models closely match the AlphaFold 3 prediction, with RMSDs of 0.568 Å and 0.563 Å relative to the AlphaFold 3 model, respectively.

(B) Using RSMD to compare the 125 AlphaFold-3 predictions for FETUA-3 with MAD-3b, MPO-1 and MDC-4 reveals a narrow distribution of RMSD for FETUA-3::MAD-3b complexes but broader distributions for FETUA-3 with MPO-1 and MDC-4. This indicates structural convergence of the similar MAD-3b structures but relatively lower structural convergence for MPO-1 and MDC-4 structures.

Figure S4 A - F

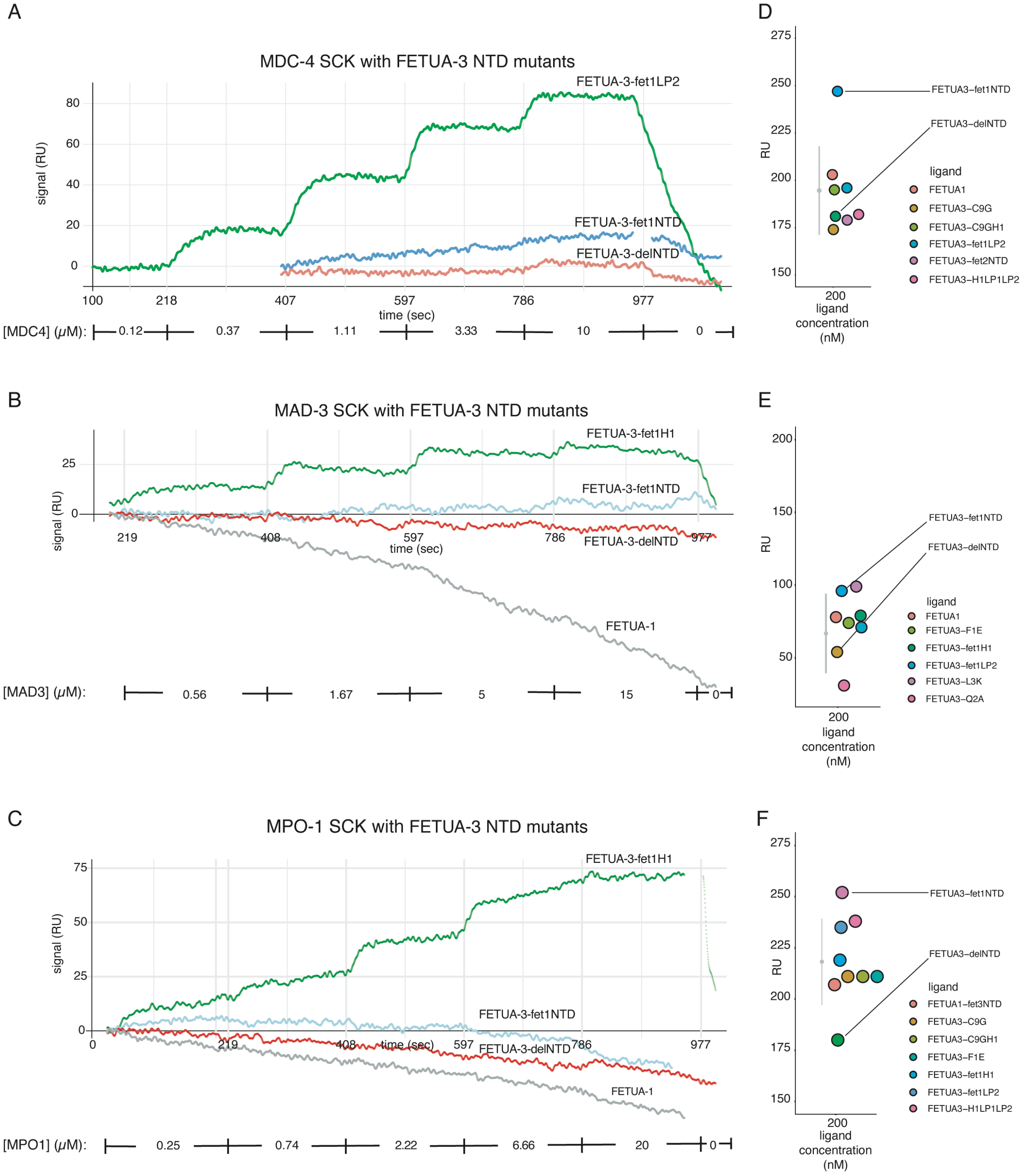


**Figure S4. The N-terminal domain of FETUA-3 is necessary for binding to MDC-4, MAD-3 and MPO-1.**

FETUA-3-delNTD protein lacks the first seven amino acids and contains an N-terminal aspartic acid. The chimeric FETUA-3-fet1NTD consists of a swap of the first seven amino acids of FETUA-3 (1-FQLAGMN-7) to the FETUA-1 sequence (1-HHLQSQI-7) while the rest of the protein is FETUA-3 sequence. FETUA-1 is a non-inhibitor of venom metalloproteinases and source of donor sequences, it lacks binding or inhibitory activity with respect to MDC-4, MAD-3 and MPO-1.

(A – C) Protein-protein interactions were measured using Surface Plasmon Resonance (SPR) in the single-cycle kinetic (SCK) format. Curves show the levels of SPR signal in response units (RU) over the duration of an experiment (seconds on x-axis). The broad concentration of enzyme applied to immobilized FETUA-3 is shown below the x-axis with brackets depicting the beginning and end of each specific concentration. Raw signal appears as a serrated line and fit data (Langmuir model with 1:1 binding) are shown as an overlayed smooth line where appropriate. For proteins without detectable binding the signal is shown relative to the signal from an internal reference measured in the same experiment.

(D - F) The signal (RU) of immobilized protein (FETUA-3 and mutants) before application of the respective analyte (MDC-4, MAD-3, MPO-1) was extracted from the raw signal files and is shown in a scatterplot with all proteins measured in the specific experiment. The mean immobilization RU is a grey point with whiskers showing the standard deviation.

(A-C) The N-terminal domain of FETUA-3 is necessary for MP binding.

(A) Binding of FETUA-3-fet1NTD and FETUA-3-delNTD to MDC-4 is undetectable. FETUA-3-fet1LP2 (for reference) binds to MDC-4 with a binding constant of ~1 µM ($K_D$ = 1.15 x $10^{-6}$ M; $B_{max}$ = 133.95, $\chi^2$ = 4.24).

(B) FETUA-3-fet1NTD and FETUA-3-delNTD binding to MAD-3 is not detected.

(C) FETUA-3-fet1NTD and FETUA-3-delNTD binding to MPO-1 is not detected. FETUA-3-fet1H1 (MAD3:H1 $K_D = 87.7 \times 10^{-6}$ M; $B_{max} = 34.55$, $\chi^2 = 4.04$; MPO1:H1 $K_D = 2.05 \times 10^{-6}$ M; $B_{max} = 76.33$, $\chi^2 = 7.85$) serves as a reference for the limit of detectable binding at these respective analyte concentration ranges.

(D-F) The immobilization levels of the FETUA proteins with modified NTDs are shown to the right (D, MDC-4; E, MAD-3; F, MPO-1).

Figure S5 A - F

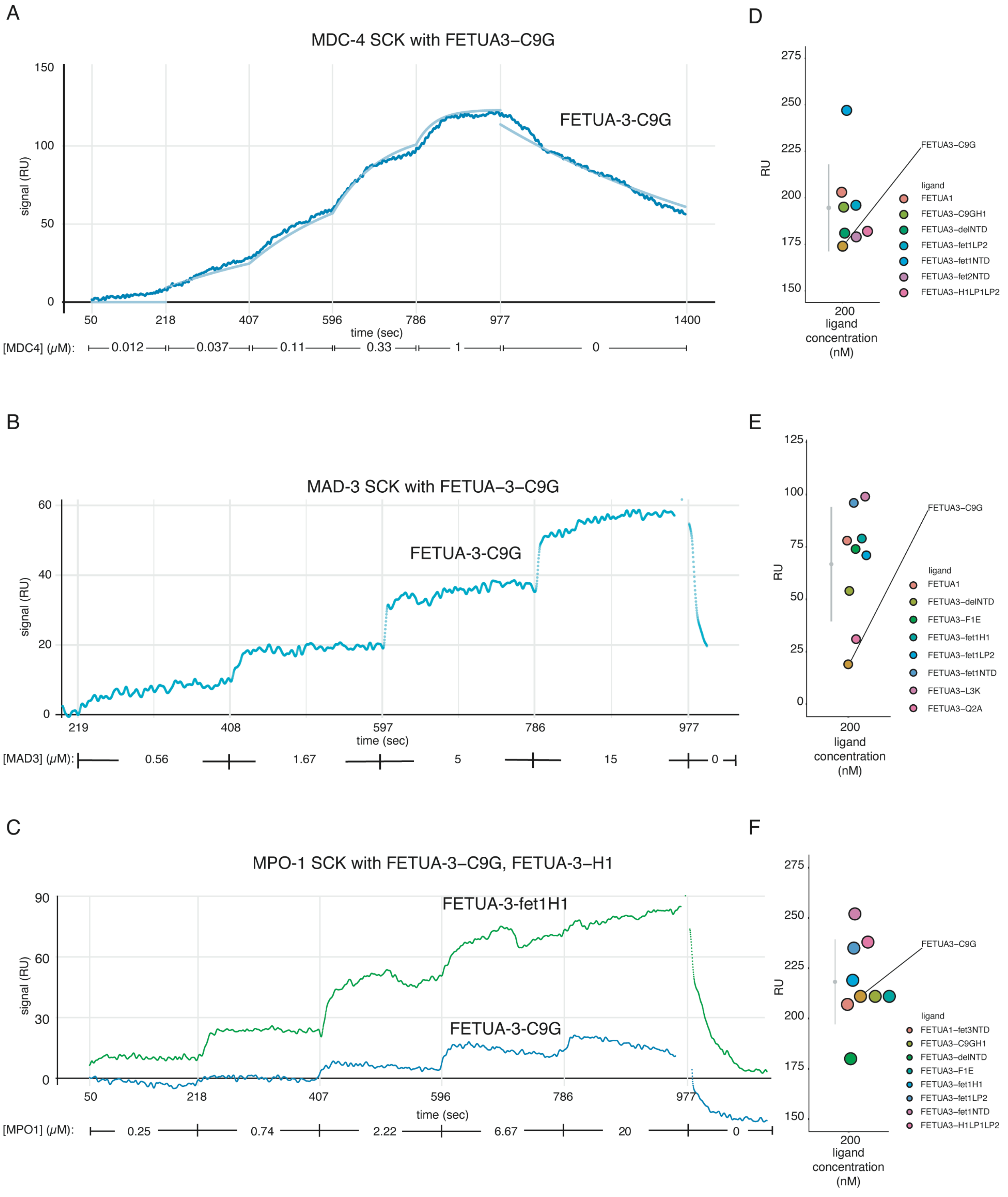


**Figure S5. The N-terminal domain of FETUA-3 is likely stabilized by a disulfide bridge between cysteines 9 (C9) and 296 (C296) which is necessary for binding to MAD-3 and MPO-1.**

(A – C) SPR binding curves as described in Figure S4.

(A) FETUA-3-C9G binds to MDC-4 with a binding constant of ~65 nM ($K_D$ = 65 x $10^{-9}$ M; $B_{max}$ = 122, $\chi^2$ = 12.13). This suggests the mutant protein is properly folded and active.

(B) An interaction between FETUA-3-C9G and MAD3 is detected but the Langmuir binding model poorly fits the data suggesting a binding constant > 88 μM (inferred from a FETUA-3-fet1H1 reference shown in Fig S6B).

(C) FETUA-3-C9G binding to MPO-1 produces only a minimal signal above background relative to FETUA-3-fet1H1 suggesting a binding constant > 2 μM (see Fig S6C).

(D – F) Scatterplots of immobilization levels of FETUA-3-C9G (D, MDC-4; E, MAD-3; F, MPO-1). See description in Figure S4.

Figure S6 A - F

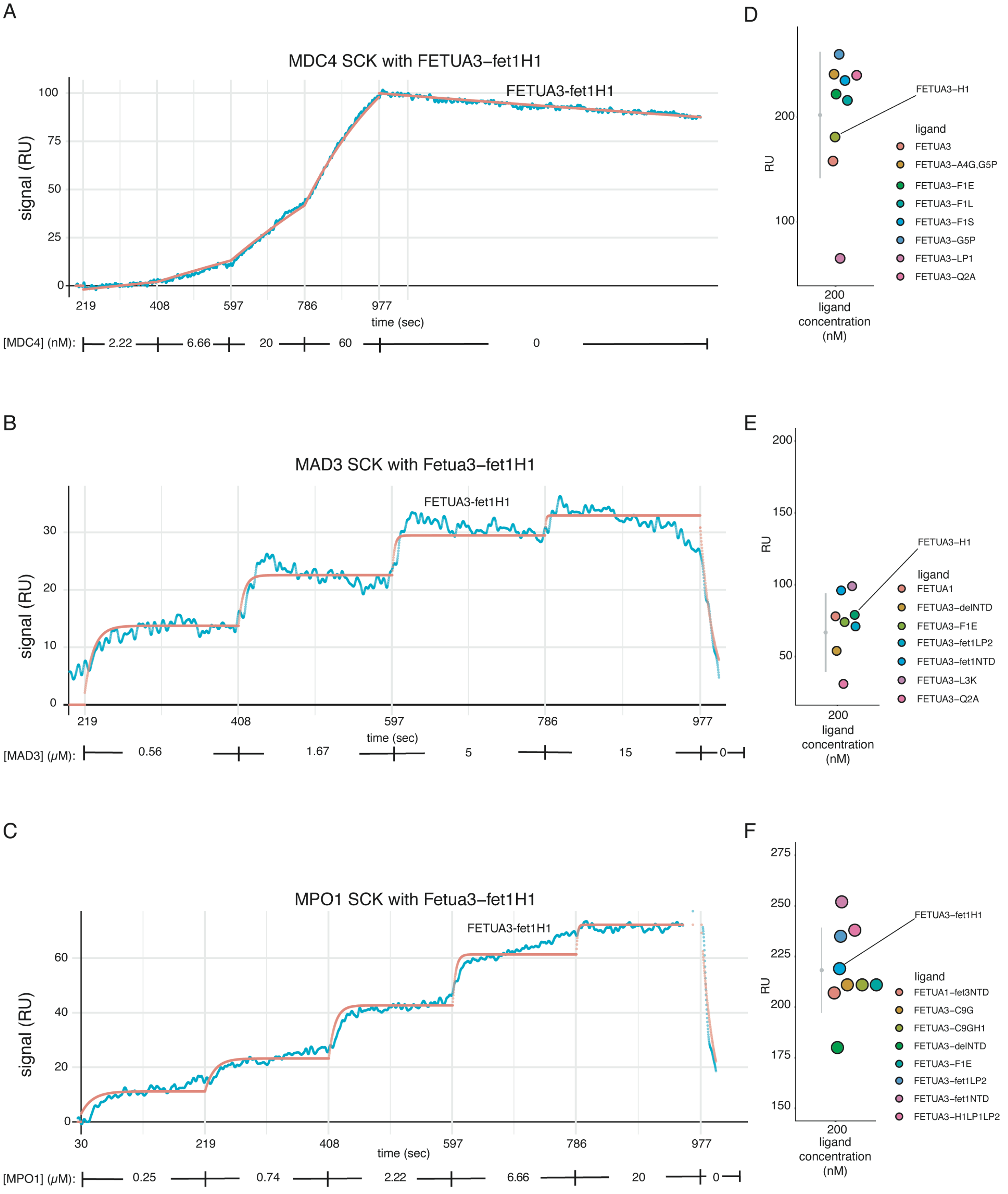


**Figure S6. A segment of alpha-helix one (H1) is necessary for high-affinity multi-target binding.**

(A – C) SPR binding curves as described in Figure S4.

(A) High affinity binding of FETUA-3-fet1H1 to MDC-4 ($K_D = 5 \times 10^{-9}$ M; $B_{max} = 171$, $\chi^2 = 2.89$) shows the chimeric protein is properly folded and active.

(B) FETUA-3-fet1H1 binding to MAD-3 is severely reduced ($K_D = 88 \times 10^{-6}$ M; $B_{max} = 35$, $\chi^2 = 4.04$).

(C) FETUA-3-fet1H1 binding to MPO-1 is severely reduced (KD = $2 \times 10^{-6}$ M; $B_{max} = 76$, $\chi^2 = 7.85$). This suggests that H1 is component of the FETUA-3::MP multi-target binding interface.

(D – F) Scatterplots of immobilization levels of FETUA-3-fet1H1 (D, MDC-4; E, MAD-3; F, MPO-1). See description in Figure S4.

Figure S7 (A-F)

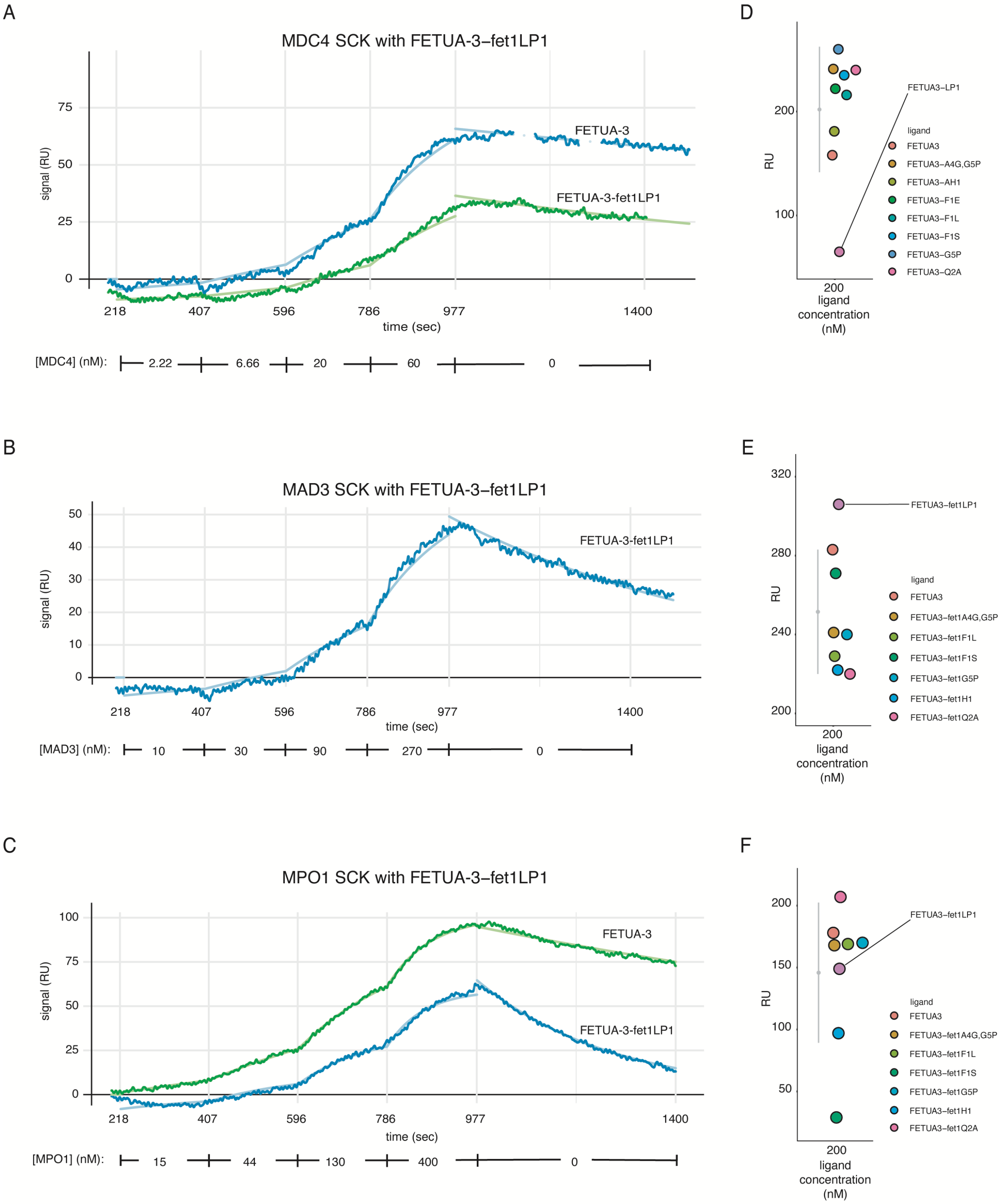


**Figure S7. Hairpin loop 1 (“Loop 1; LP1”) has modest effects on multi-target binding.**

(A – C) SPR binding curves as described in Figure S4.

(A) FETUA-3-fet1LP1 binding to MDC-4 is reduced 5-fold (increased $K_D$).

(B) FETUA-3-fet1LP1 binding to MAD-3 is reduced 8-fold (increased $K_D$).

(C) FETUA-3-fet1LP1 binding to MPO-1 is reduced 7-fold (increased $K_D$). The modest reduction in binding of FETUA-3-fet1LP1 to MPs appears to be tolerated with respect to inhibition because only a slight loss reduction of inhibition is detected for FETUA-3-fet1LP1.

(D – F) Scatterplots of immobilization levels of FETUA-3-fet1H1 (D, MDC-4; E, MAD-3; F, MPO-1). See description in Figure S4.

Figure S8 A - F

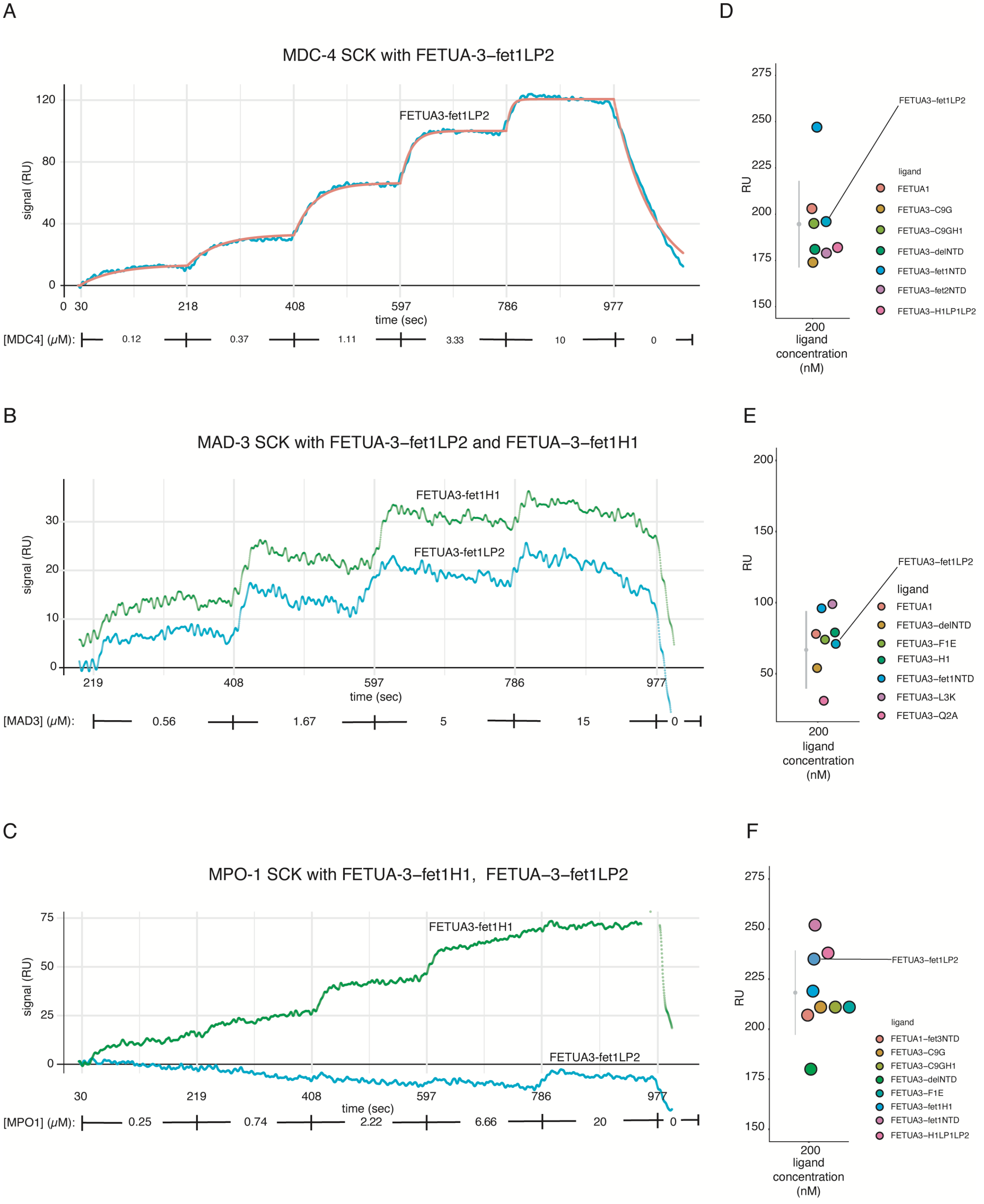

A
MDC-4 SCK with FETUA-3–fet1LP2
FETUA3-fet1LP2
signal (RU)
120
80
40
0
0 30
218
408
597
786
977
time (sec)
[MDC4] (μM):
0.12
0.37
1.11
3.33
10
0
B
MAD-3 SCK with FETUA-3–fet1LP2 and FETUA–3–fet1H1
FETUA3-fet1H1
FETUA3-fet1LP2
signal (RU)
30
20
10
0
219
408
597
786
977
time (sec)
[MAD3] (μM):
0.56
1.67
5
15
0
C
MPO-1 SCK with FETUA-3–fet1H1, FETUA–3–fet1LP2
FETUA3-fet1H1
FETUA3-fet1LP2
signal (RU)
75
50
25
0
30
219
408
597
786
977
time (sec)
[MPO1] (μM):
0.25
0.74
2.22
6.66
20
0
D
RU
275
250
225
200
175
150
200
ligand
concentration
(nM)
FETUA3–fet1LP2
ligand
FETUA1
FETUA3–C9G
FETUA3–C9GH1
FETUA3–delNTD
FETUA3–fet1NTD
FETUA3–fet2NTD
FETUA3–H1LP1LP2
E
RU
200
150
100
50
200
ligand
concentration
(nM)
FETUA3–fet1LP2
ligand
FETUA1
FETUA3–delNTD
FETUA3–F1E
FETUA3–H1
FETUA3–fet1NTD
FETUA3–L3K
FETUA3–Q2A
F
RU
275
250
225
200
175
150
200
ligand
concentration
(nM)
FETUA3–fet1LP2
ligand
FETUA1–fet3NTD
FETUA3–C9G
FETUA3–C9GH1
FETUA3–delNTD
FETUA3–F1E
FETUA3–fet1H1
FETUA3–fet1NTD
FETUA3–H1LP1LP2

**Figure S8. Hairpin loop 2 ("Loop 2; LP2") is necessary for high-affinity multi-target binding.**

(A – C) SPR binding curves as described in Figure S4.

(A) FETUA-3-fet1LP2 binding to MDC4 is severely reduced with a ~344-fold decrease in the dissociation constant ($K_D$ = 1.15 x $10^{-6}$ M; $B_{max}$ = 134, $\chi^2$ = 4.24). The reduced binding constant relative to FETUA-3 is partly due to a 6-fold decrease in on-rate but mainly is the result of a 58-fold faster off-rate which suggests a role for loop2 in stabilization of the FETUA-3::MP inhibition complex. The diminished but reproducible binding of FETUA-3-fet1LP2 to MDC-4 is important because it establishes a limit for the detection of weak binding thus allowing us to infer that binding of MAD-3 and MPO-1 to FETUA-3-fet1LP2 is "weak" with a binding constant greater than 1 μM.

(B) FETUA-3-fet1LP2 binding to MAD-3 is weak. FETUA-3-fet1H1 is an internal experimental reference that bound ($K_D$ = 88 x $10^{-6}$ M) in the experiment at high concentrations of MAD-3.

(C) FETUA-3-fet1LP2 binding to MPO-1 is weak. FETUA-3-fet1H1 again is a reference that bound ($K_D$ = 2 x $10^{-6}$ M) in the experiment at high concentrations of MPO-1.

(D – F) Scatterplots of immobilization levels of FETUA-3-fet1H1 (D, MDC-4; E, MAD-3; F, MPO-1). See description in Figure S4.

Figure S9 A - F

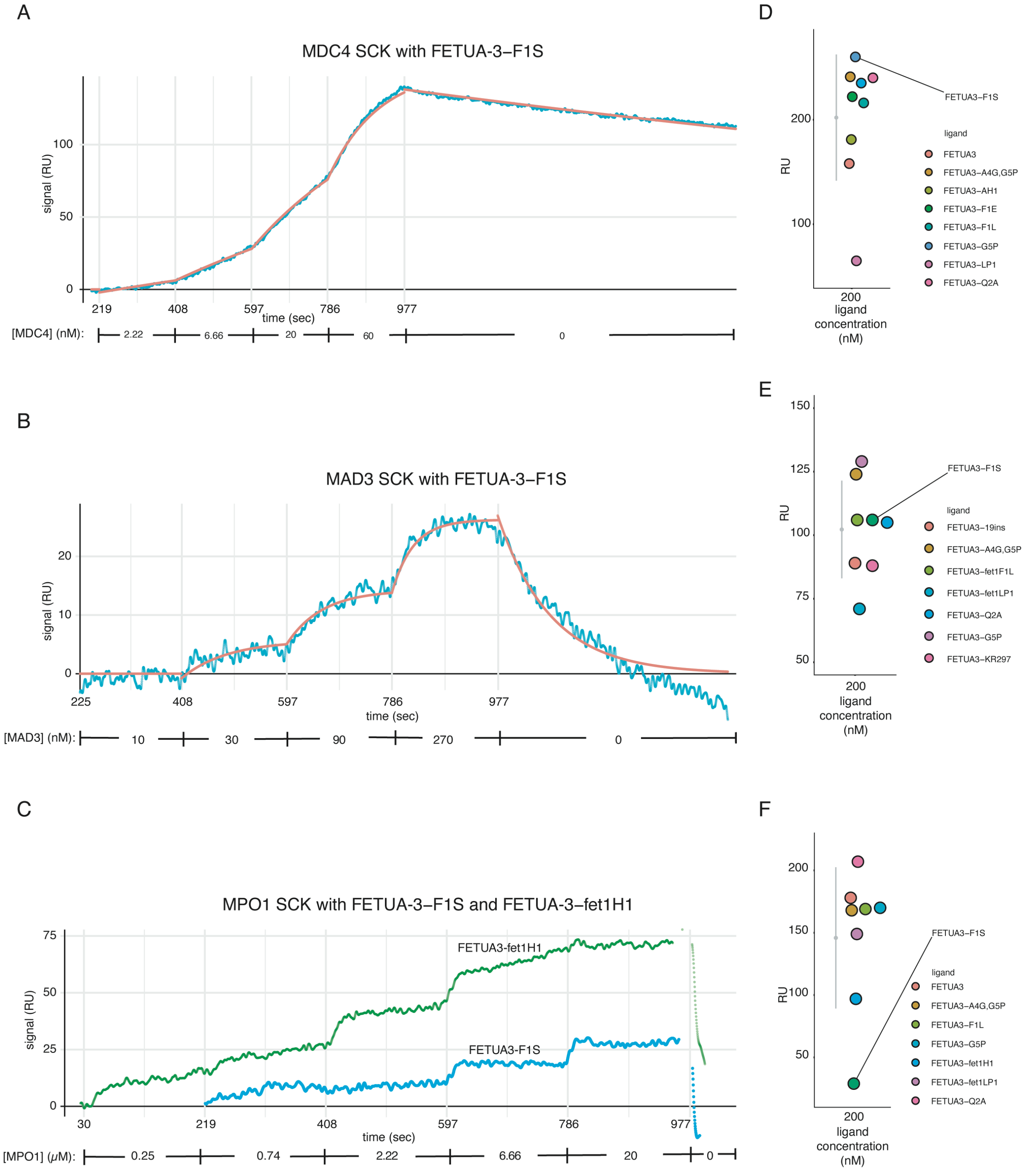


**Figure S9. The N-terminal phenylalanine (F1) of FETUA-3 is necessary for multi-target binding.**

(A – C) SPR binding curves as described in Figure S4. Substitution of Phe-1 with a serine residue (FETUA-3-F1S) differentially affects binding to MDC-4, MAD-3 and MPO-1.

(A) FETUA-3-F1S binding to MDC-4 is similar (~1.2-fold reduction) to FETUA-3 ($K_D$: WT = $3.44 \times 10^{-9}$ M, F1S = $4.03 \times 10^{-9}$ M).

(B) FETUA-3-F1S binding to MAD-3 is reduced ~19-fold ($K_D$: WT = $1.3 \times 10^{-8}$ M, F1S = $24.5 \times 10^{-8}$ M).

(C) FETUA-3-F1S binding to MPO-1 is weak ($K_D$: H1 (reference) = $2 \times 10^{-6}$ M; F1S > $2 \times 10^{-6}$ M).

(D – F) Scatterplots of immobilization levels of FETUA-3-F1S (D, MDC-4; E, MAD-3; F, MPO-1). See description in Figure S4.

Figure S10 A - F

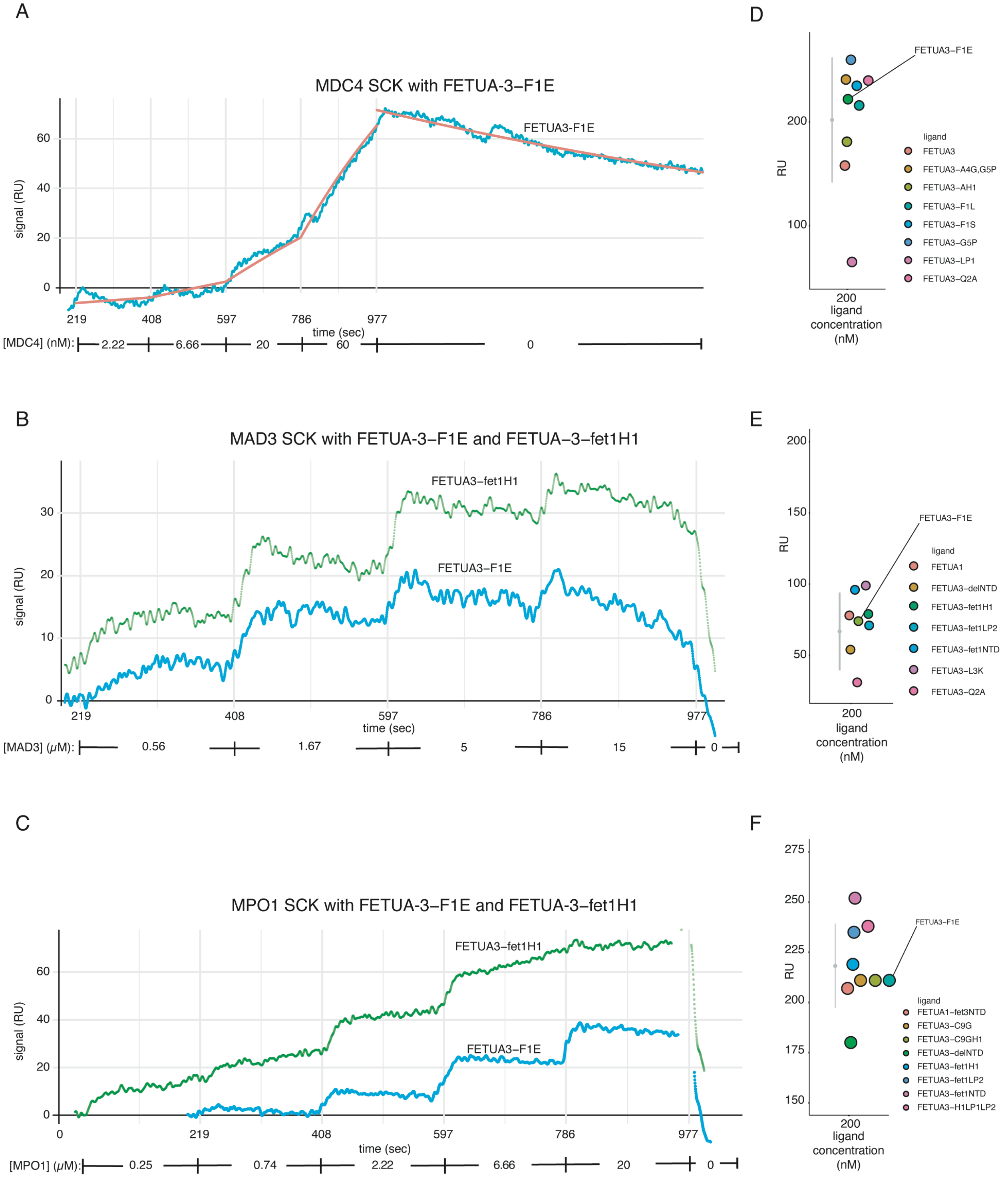


**Figure S10**. **The N-terminal phenylalanine (F1) of FETUA-3 is necessary for multi-target binding.**

(A – C) SPR binding curves as described in Figure S4. Substitution of Phe-1 with a glutamic acid residue (FETUA-3-F1E) differentially affects binding to MDC-4, MAD-3 and MPO-1.

(A) FETUA-3-F1E binding to MDC-4 is decreased five-fold ($K_D$: WT = 3.44 x $10^{-9}$ M, F1E = 19.5 x $10^{-9}$ M).

(B) FETUA-3-F1E binding to MAD-3 is not detected ($K_D$: H1 (reference) = 88 x $10^{-6}$ M, F1E > 88 x $10^{-6}$ M).

(C) FETUA-3-F1E binding to MPO-1 is not detected ($K_D$: H1 (reference) = 2 x $10^{-6}$ M, F1E > 2 x $10^{-6}$ M).

(D – F) Scatterplots of immobilization levels of FETUA-3-F1E (D, MDC-4; E, MAD-3; F, MPO-1). See description in Figure S4.

Figure S11

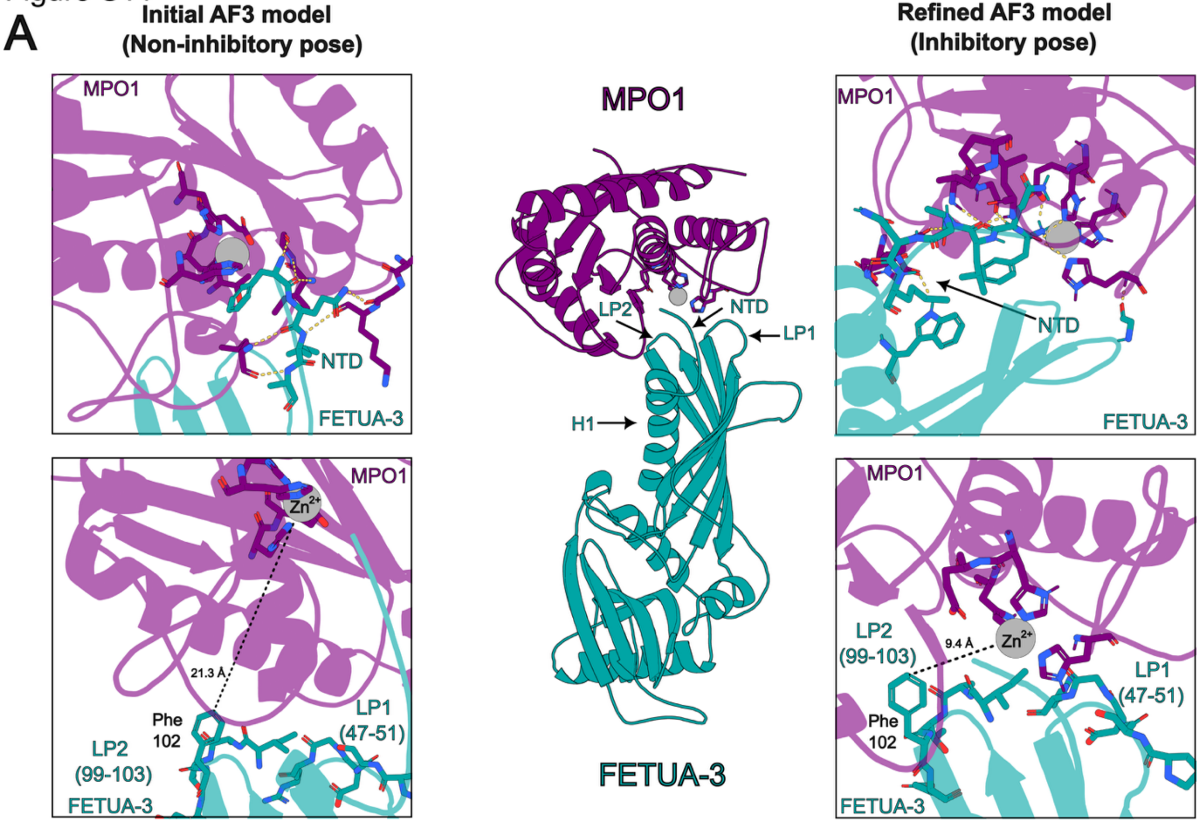
A
Initial AF3 model
(Non-inhibitory pose)
Refined AF3 model
(Inhibitory pose)
MPO1
NTD
FETUA-3
MPO1
LP2
NTD
LP1
H1
MPO1
NTD
FETUA-3
MPO1
Zn2+
21.3 Å
Phe
102
LP2
(99-103)
LP1
(47-51)
FETUA-3
FETUA-3
MPO1
LP2
(99-103)
9.4 Å
Zn2+
LP1
(47-51)
Phe
102
FETUA-3

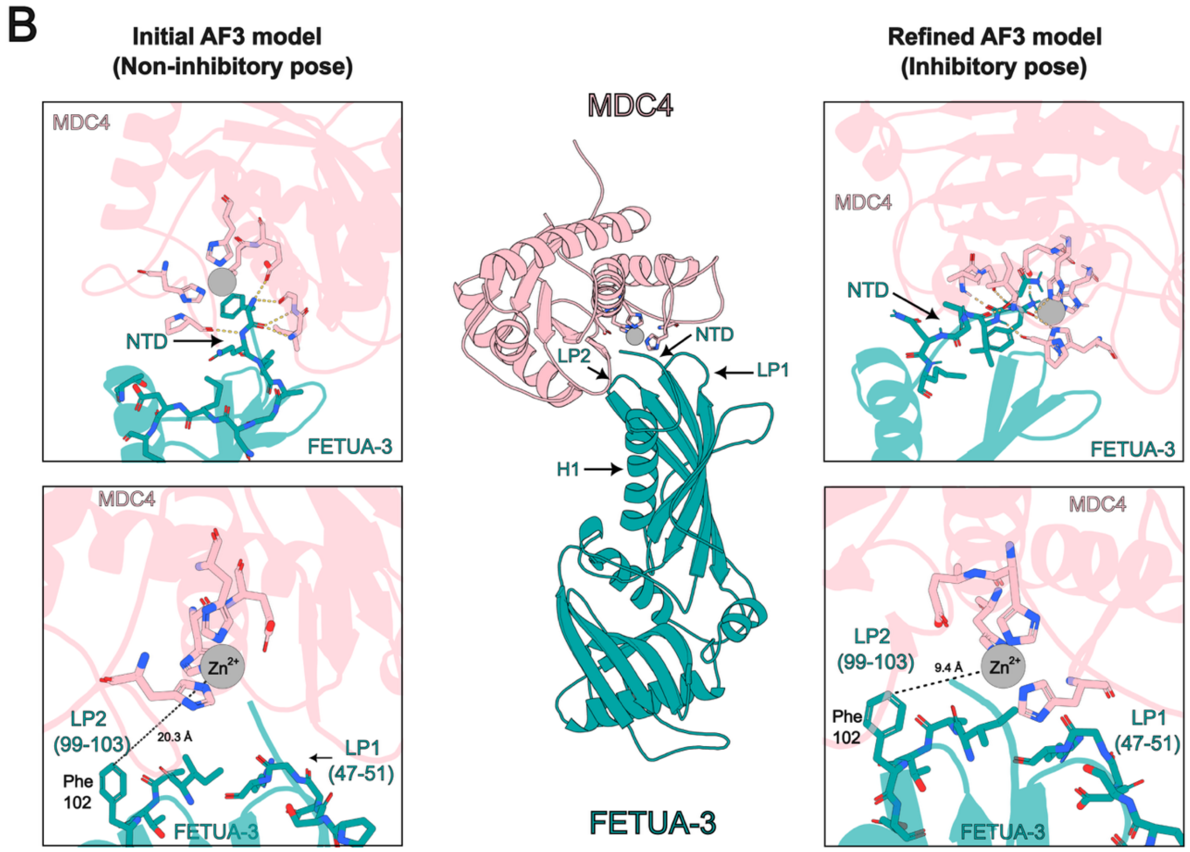
B
Initial AF3 model
(Non-inhibitory pose)
Refined AF3 model
(Inhibitory pose)
MDC4
NTD
FETUA-3
MDC4
LP2
NTD
LP1
H1
MDC4
NTD
FETUA-3
MDC4
Zn2+
LP2
(99-103)
20.3 Å
Phe
102
LP1
(47-51)
FETUA-3
FETUA-3
MDC4
LP2
(99-103)
9.4 Å
Zn2+
Phe
102
LP1
(47-51)
FETUA-3

**Figure S11. Initial and refined AlphaFold3 predictions of FETUA-3 in complex with MPO-1 and MDC-4.**

**(A)** Predicted structure of the FETUA-3–MPO1 complex. Top left, the initial AlphaFold3 (AF3) model positions the N-terminal domain of FETUA-3 in a non-inhibitory orientation, forming limited electrostatic interactions with regions surrounding the MPO1 active site. Top right, the refined AF3 model repositions the N-terminal domain into an inhibitory pose resembling that observed in the FETUA-3–MAD3B complex, with extensive electrostatic interactions both proximal and distal to the active site. Bottom left, in the initial model, loop 1 (LP1) and loop 2 (LP2) are positioned distant from the enzyme, incompatible with inhibition. The position of Phe102 is shown as a reference for inhibitor–enzyme proximity, with a measured distance of 21.3 Å. Bottom right, in the refined model, LP1 and LP2 are positioned adjacent to the enzyme in a conformation compatible with inhibition, reducing the corresponding distance to 9.4 Å.

**(B)** Predicted structure of the FETUA-3–MDC4 complex. Top left, the initial AF3 model places the N-terminal domain of FETUA-3 in a non-inhibitory orientation with limited electrostatic interactions near the MDC4 active site. Top right, the refined AF3 model positions the N-terminal domain in an inhibitory pose similar to that observed for MAD3B, forming extensive electrostatic interactions with regions proximal and distal to the active site. Bottom left, the initial model predicts LP1 and LP2 distant from the enzyme, inconsistent with an inhibitory conformation. Phe102 is highlighted as a positional reference, showing a distance of 20.3 Å relative to MDC4. Bottom right, the refined model predicts LP1 and LP2 are adjacent to the enzyme in a conformation compatible with inhibition, decreasing the corresponding distance to 9.4 Å.

Figure S12 Rosetta Interface analysis of original and refined models

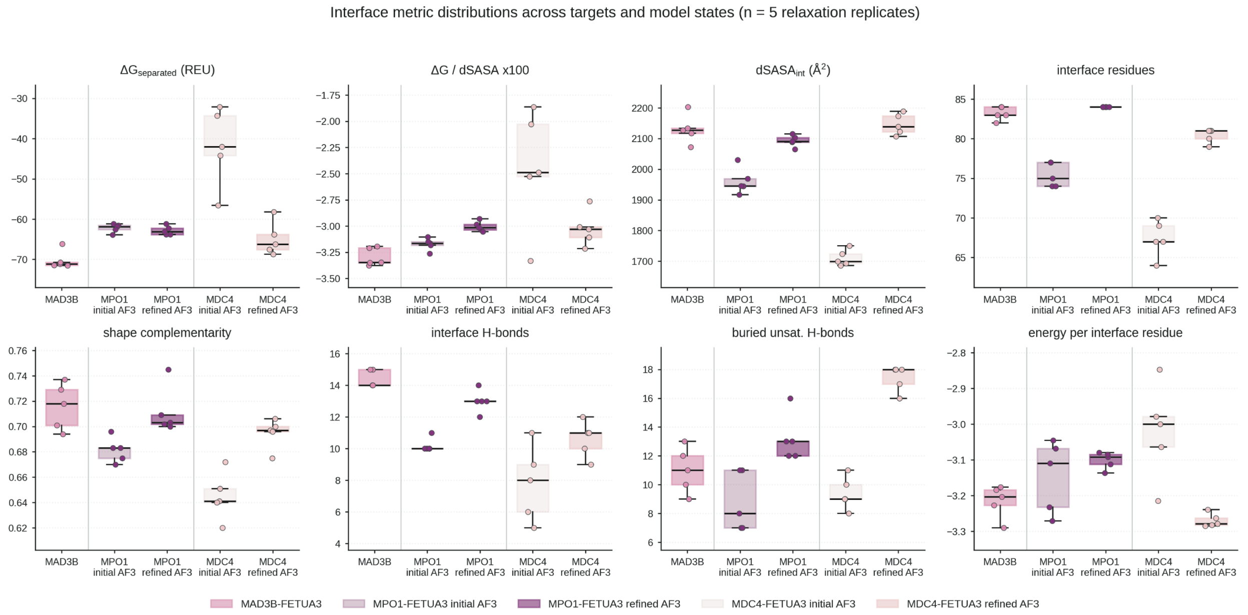


**Figure S12. Evaluation of initial and refined models by Rosetta InterfaceAnalyzer.** AF3 predicted MAD-3b–FETUA3 with high confidence and a single consistent binding mode, whereas the MPO-1–FETUA3 and MDC-4–FETUA3 predictions were low-confidence and yielded poses inconsistent with an inhibitory binding mode. These two complexes were therefore manually re-docked to match the MAD3B–FETUA3 reference and re-scored. Each model state was subjected to Rosetta FastRelax and its interface analyzed; distributions show n = 5 relaxation replicates (points), with boxes spanning the interquartile range (IQR), median line, and whiskers extending to 1.5 × IQR. Gray lines separate targets; colors denote model state (key). Panels: ΔG_separated; ΔG normalized to buried area; dSASA_int (Å²); interface residue count; shape complementarity (Sc); cross-interface hydrogen bonds; buried unsatisfied hydrogen bonds; and ΔG per interface residue.

Figure S13 Distribution of distances between backbone elements of FETUA-3-Phe1 and the zinc ion in the MAD-3b structures. Stability of complexes of FETUA-3::MAD-3b, ::MPO-1 and ::MDC-4 during QM/MM and MM Molecular Dynamic simulations.

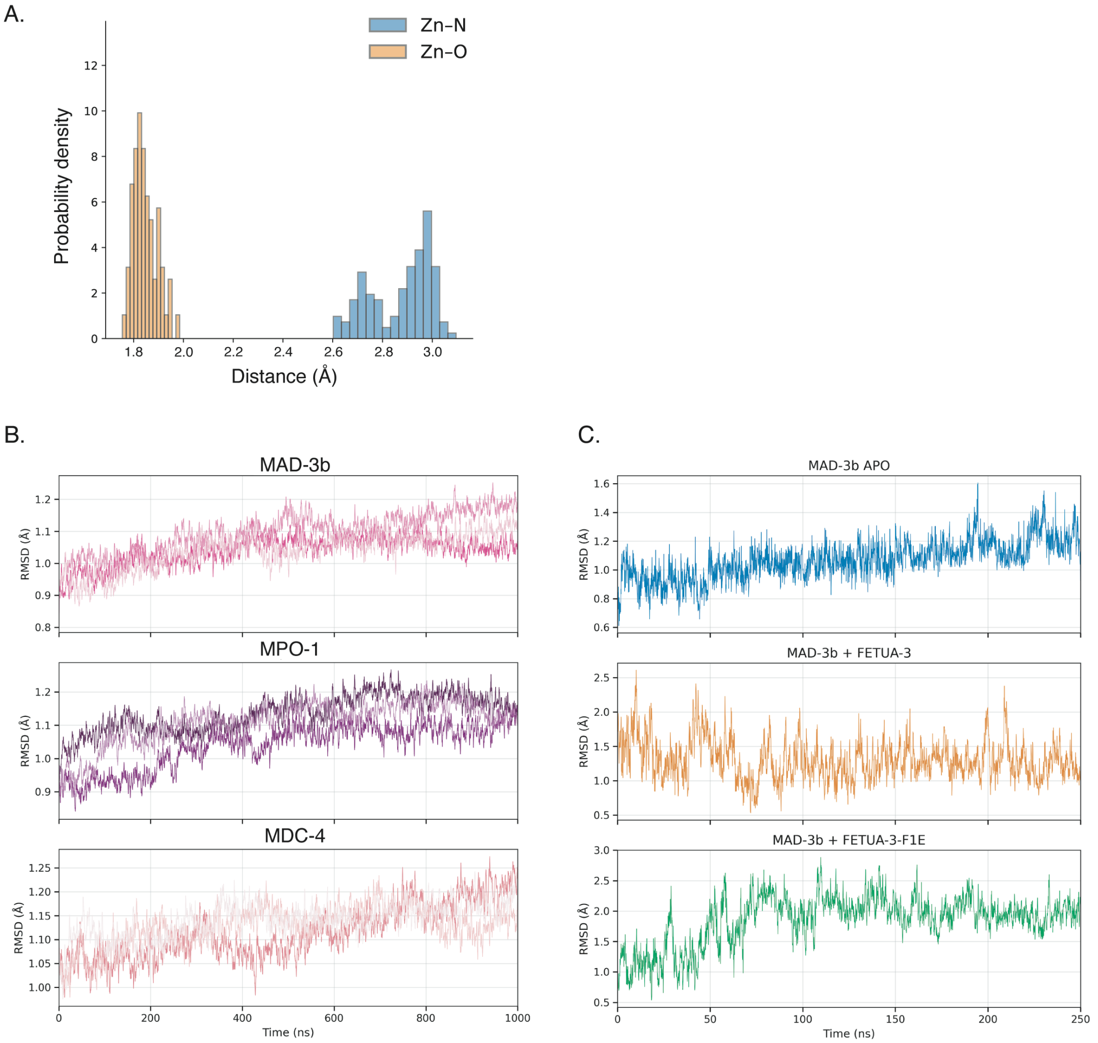


**Figure S13. Distribution of distances between backbone elements of FETUA-3-Phe1 and the $Zn^{2+}$ in the MAD-3b models. Stability of complexes of FETUA-3::MAD-3b, ::MPO-1 and ::MDC-4 during QM/MM and MM Molecular Dynamic simulations.**

(A) The distance (Å) between the active site $Zn^{2+}$ (Zn) and backbone elements of FETUA-3-Phe1 (N-terminal amino nitrogen (N) and carbonyl oxygen (O)) for 125 MAD-3b::FETUA-3 AF3 models. The mean Zn-O (tan) distance (1.8 ± 0.05) is less than the mean Zn-N (blue) distance (2.9 ± 0.2 Å). This suggests the carbonyl oxygen may directly coordinate $Zn^{2+}$ and does not support direct coordination of $Zn^{2+}$ via the N-terminal amine.

(B) Root-mean-square deviation (RMSD) of the predicted complexes throughout the quantum mechanics/molecular mechanics (QM/MM) molecular dynamics (MD) simulations for MAD-3b (top panel), MPO-1 (middle), MDC-4(bottom). Three independent replicates are shown in different colors in each panel.

(C) Root-mean-square deviation (RMSD) of the structures throughout the production trajectories of the classical molecular mechanics (MM) molecular dynamics (MD) simulations for APO MAD-3b (top), MAD-3b::FETUA-3 complex (middle) and MAD-3b::FETUA-3-F1E (bottom). The initial 50 nanoseconds (ns) of each trajectory were considered equilibration while the remainder was used for analysis. The color scheme is the same used in Figure 6D.

Figure S14 A - F

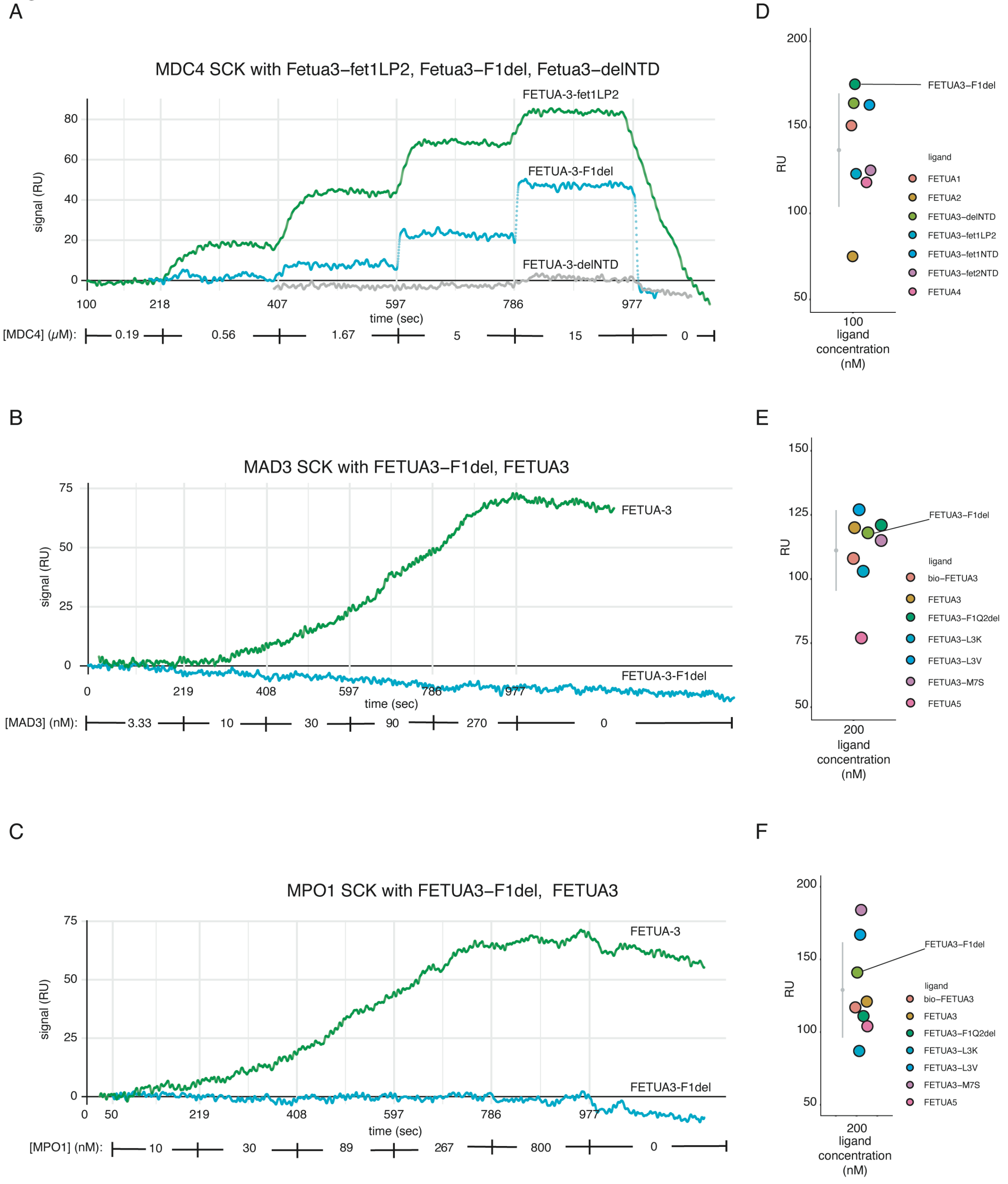


**Figure S14. The N-terminal phenylalanine (F1) of FETUA-3 is necessary for multi-target binding.**

(A – C) SPR binding curves as described in Figure S4. Deletion of Phe-1 (FETUA-3-F1del) severely reduces or abolishes binding to MDC-4, MAD-3 and MPO-1.

(A) FETUA-3-F1del binding to MDC-4 is severely reduced (~465-fold reduction of the dissociation constant; $K_D$: F1del = 16.7 x $10^{-6}$ M, reference LP2 = 1.15 x $10^{-6}$ M). The undetectable binding of FETUA-3-delNTD (first seven amino acids deleted from the N-terminus; grey line) is shown for comparison with the weak but detectable binding of FETUA-3-F1del.

(B) FETUA-3-F1del binding to MAD-3 is not detected ($K_D$: F1del > $10^{-6}$ M, reference H1 = 88 x $10^{-6}$ M).

(C) FETUA-3-F1del binding to MPO-1 is not detected ($K_D$: F1del > $10^{-6}$ M, reference WT = 16.7 x $10^{-9}$ M).

(D – F) Scatterplots of immobilization levels of FETUA-3-F1del (D, MDC-4; E, MAD-3; F, MPO-1). See description in Figure S4.

Figure S15 Structure predicitions identify key interface contacts between MAD-3b and FETUA-3 NTD residues Q2 and L3.

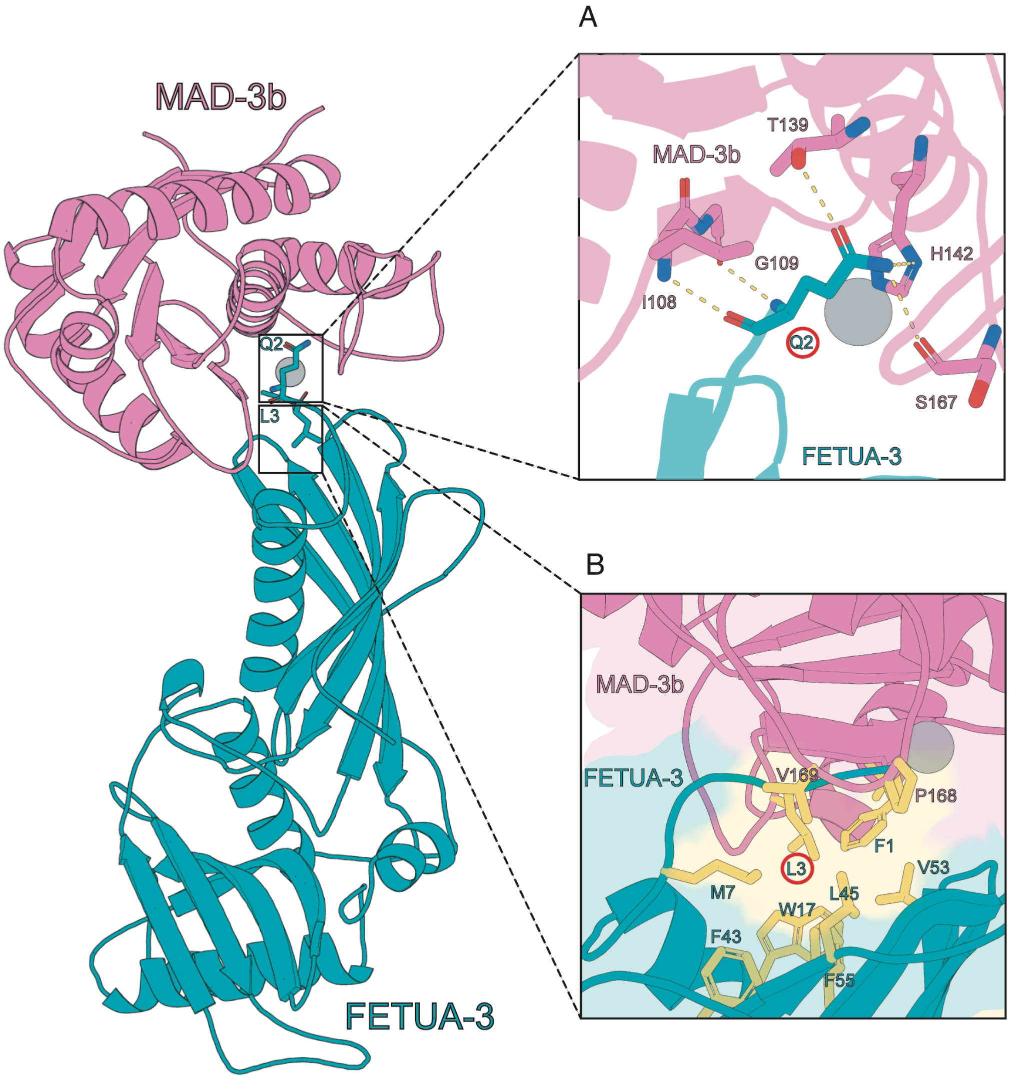


**Figure S15. Structure predictions identify putative contacts at the FETUA-3::MAD-3b interface.** (A) FETUA-3 Gln2 may contact Ile-108 of MAD-3b and (B) FETUA-3 Leu3 is in a hydrophobic pocket of MAD3b.

Figure S16 A - F

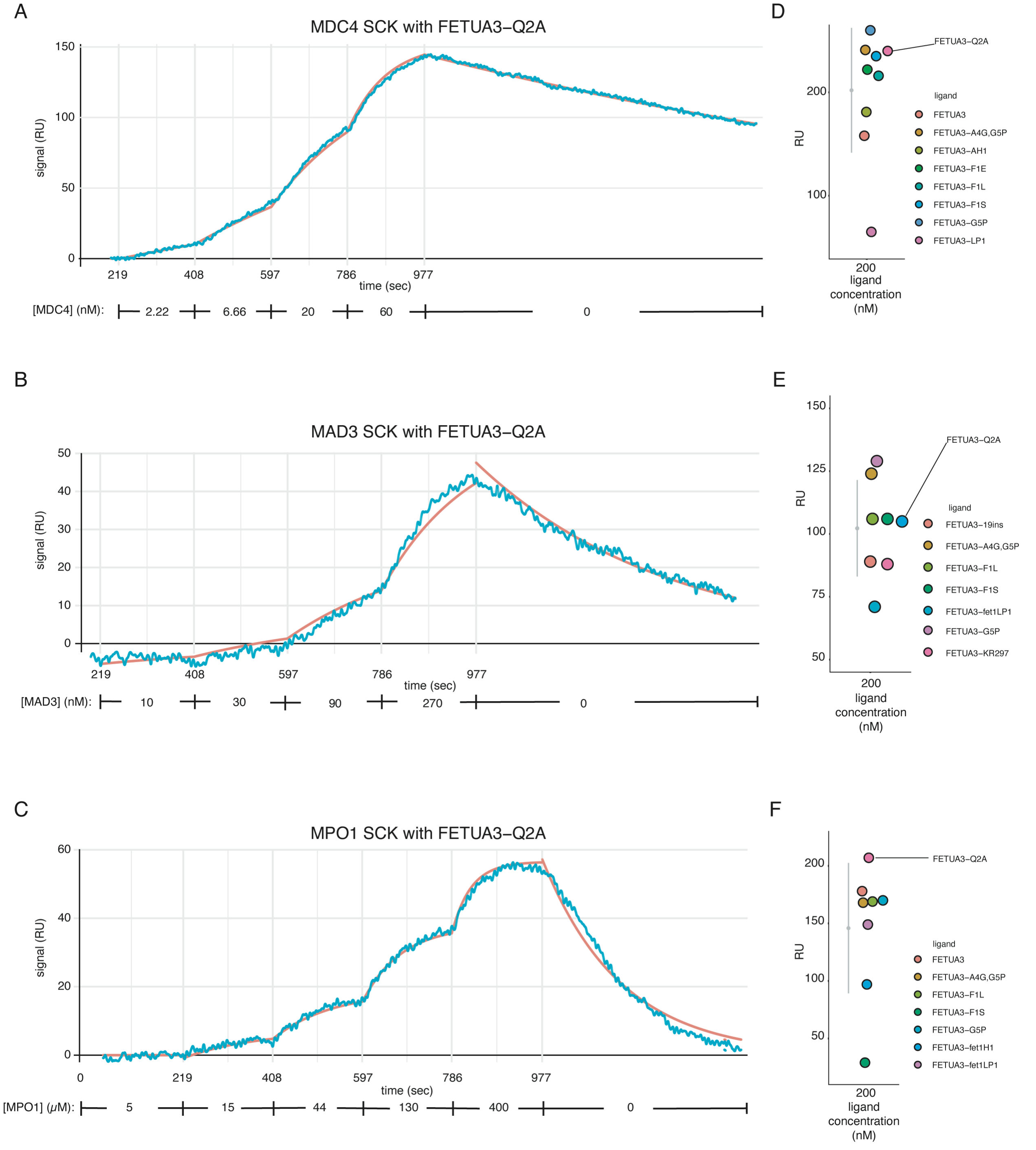


**Figure S16. Glutamine in the second position (Gln2, Q2) of the FETUA-3 NTD stabilizes the FETUA-3::MP complex.**

(A – C) SPR binding curves as described in Figure S4. FETUA-3-Q2A on-rates were like those measured for FETUA-3 however increased off-rates were detected for MDC-4, MAD-3 and

MPO-1. This suggests that glutamine (Q2) may be necessary to stabilize the inhibitor-enzyme complex for all MP targets.

(A) FETUA-3-Q2A binds to MDC-4 with a similar on-rate to that of FETUA-3. ($k_{on}$: Q2A = 17 x $10^4$, WT = 7 x $10^4$) but with a slight (1.9-fold) increase in off-rate.

(B) FETUA-3-Q2A binds to MAD-3 with a similar on-rate to that of FETUA-3 ($k_{on}$: Q2A = 2.97 x $10^4$, WT = 2.84 x $10^4$) but with an increased (6.9-fold) off-rate.

(C) FETUA-3-Q2A binds to MPO-1 with a similar on-rate to that of FETUA-3 ($k_{on}$: Q2A = 4.7 x $10^4$, WT = 2.6 x $10^4$ $M^{-1}s^{-1}$) but with an increased (9.4-fold) off-rate.

(D – F) Scatterplots of immobilization levels of FETUA-3-Q2A (D, MDC-4; E, MAD-3; F, MPO-1). See description in Figure S4.

Figure S17 A - F

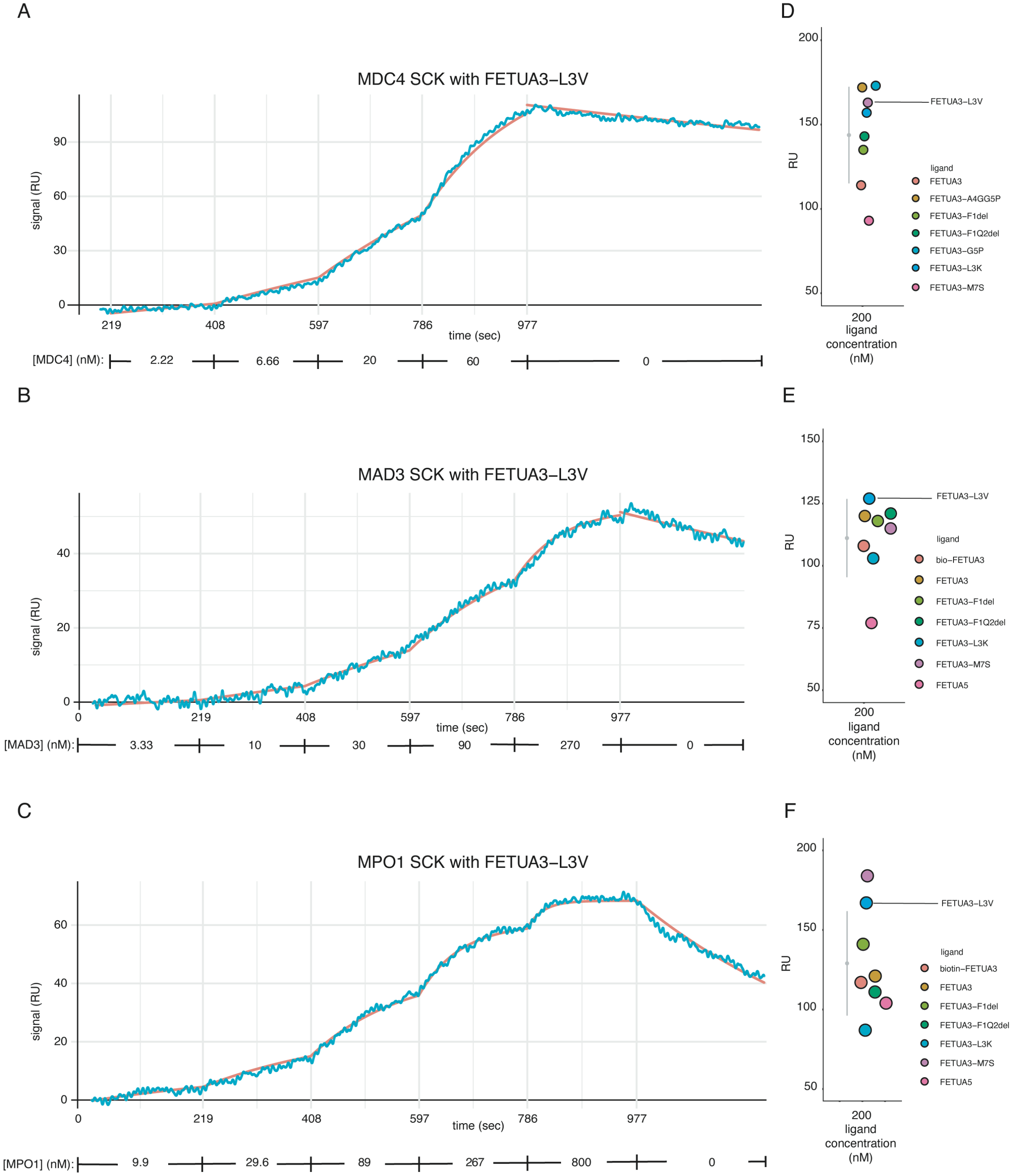


**Figure S17. Amino acid substitution of leucine-3 with valine (FETUA-3-L3V) does not disrupt binding to MDC-4, MAD-3 and MPO-1.**

(A – C) SPR binding curves as described in Figure S4. Substitution of leucine-3 to a valine has minimal effect on binding to MDC-4, MAD-3 and MPO-1.

(A) FETUA-3-L3V binding to MDC-4 is similar to FETUA-3 (1.03-fold increase in $K_D$).

(B) FETUA-3-L3V binding to MAD-3 is similar to FETUA-3 (1.36-fold increase in $K_D$).

(C) FETUA-3-L3V binding to MPO-1 is similar to FETUA-3 (2.2-fold increase in $K_D$).

(D – F) Scatterplots of immobilization levels of FETUA-3-L3V (D, MDC-4; E, MAD-3; F, MPO-1). See description in Figure S4.

Figure S18 A - F

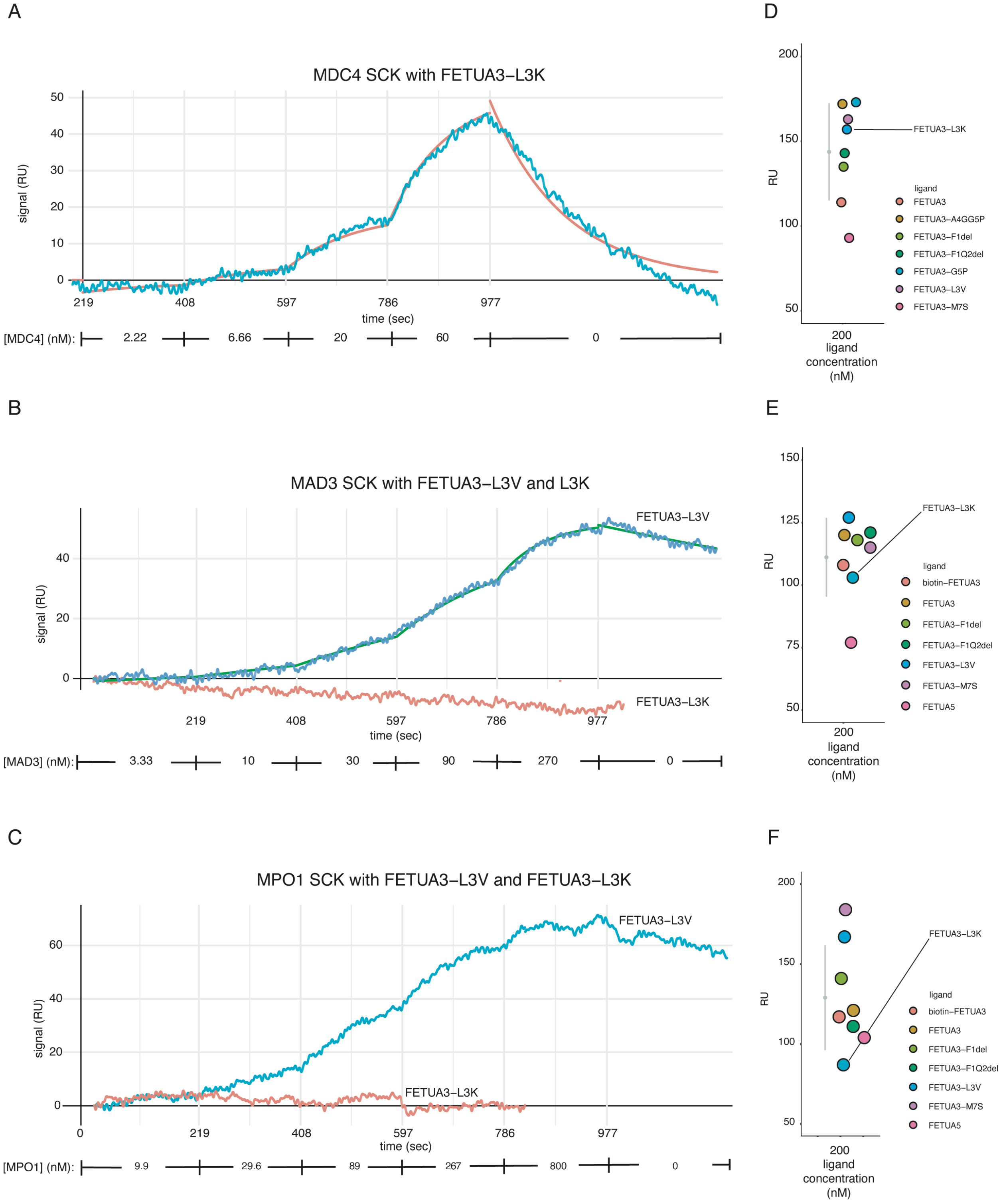


**Figure S18. Amino acid substitution of leucine-3 with lysine (FETUA-3-L3K) severely disrupts binding to MDC-4, MAD-3 and MPO-1.**

(A – C) SPR binding curves as described in Figure S4.

(A) FETUA-3-L3K binding to MDC-4 is reduced ~57-fold ($K_D$: L3K = 1.95 x $10^{-7}$ M, WT = 3.4 x $10^{-9}$ M). The increase of the L3K::MDC4 binding constant is primarily attributable to a faster off rate suggesting that L3 of the NTD likely functions in stabilizing the FETUA-3::MP inhibitory complex.

(B) FETUA-3-L3K binding to MAD-3 is not detected (L3V shown as internal control).

(C) FETUA-3-L3K binding to MPO-1 is not detected (L3V shown as internal control).

(D – F) Scatterplots of immobilization levels of FETUA-3-L3K (D, MDC-4; E, MAD-3; F, MPO-1). See description in Figure S4.
